# A two-step mechanism for RIG-I activation by influenza virus mvRNAs

**DOI:** 10.1101/2025.01.04.630666

**Authors:** Emmanuelle Pitré, Karishma Bisht, Kaleigh A. Remick, Amir Ghorbani, Jonathan W. Yewdell, Elizaveta Elshina, Aartjan J.W. te Velthuis

**Affiliations:** Lewis Thomas Laboratory, Department of Molecular Biology, Princeton University, 08544 New Jersey, United States; Department of Pathology, Addenbrooke’s Hospital, University of Cambridge, Cambridge CB2 2QQ, United Kingdom; Cellular Biology Section, Laboratory of Viral Diseases, National Institute of Allergy and Infectious Diseases, Bethesda, MD 20892, USA

**Keywords:** influenza A virus, transient RNA structure, RNA polymerase, RIG-I, interferon, mvRNA, cmvRNA, cvRNA, ccRNA

## Abstract

Influenza A virus (IAV) non-canonical replication products can be bound by host pathogen sensors, such as retinoic acid-inducible gene I (RIG-I). However, innate immune activation is infrequent in cell culture infection, in particular by adapted strains. Moreover, it is not understood why non-canonical IAV RNAs activate RIG-I in a sequence- or RNA structure-dependent manner. We therefore hypothesized that multiple errors need to occur before influenza virus RNA synthesis activates innate immune signaling. To test this idea, we investigated whether RIG-I activation is stimulated by the non-canonical or aberrant transcription of mini viral RNAs (mvRNA), a <125 nt long RNA that is overexpressed in pandemic and highly pathogenic IAV infections. Using mvRNA sequences identified in tissue culture and ferret infections, we find that mvRNAs can cause non-canonical transcription termination through a truncated 5ʹ polyadenylation signal or a 5ʹ transient RNA structure that interrupts polyadenylation. The resulting capped complementary RNAs (ccRNA) can stimulate the release of a template mvRNA in vitro. Finally, we find that both mvRNA and ccRNA sequences can be bound by RIG-I in cell culture and that blocking mvRNA transcription with baloxavir reduces IFN promoter activity. Overall, our findings indicate that sequential rounds of non-canonical or aberrant viral replication and transcription are needed before mvRNAs trigger innate immune signaling in a sequence-dependent manner.

## Introduction

Influenza A viruses (IAV) cause moderate to severe respiratory disease in humans. Activation and dysregulation of the innate immune system play an important role in the outcome of IAV infection (1–3). Various viral factors affect innate immune dysregulation, including the RNAs that IAVs produce (1, 4, 5). Binding of IAV RNAs to host pathogen receptor retinoic acid-inducible gene I (RIG-I) is assumed to be the onset of the response to IAV infection (6–8), although other host proteins, such as interferon (IFN)-gamma inducible protein 16 (IFI16), bind IAV RNAs as well (9). Once activated, RIG-I triggers the expression of interferon and proinflammatory genes, and a dysregulation of their function can impact disease severity (3, 10, 11). Variation has been observed among non-canonical RNAs in their ability to activate RIG-I that is independent of RNA length (12–14). Presently, the molecular steps that lead from IAV RNA synthesis to IAV RNA binding by RIG-I are not understood.

The IAV genome consists of 8 segments of single-stranded, negative sense RNA (vRNA) that contain 5ʹ triphosphorylated, partially complementary 5ʹ and 3ʹ promoter termini (15–17). The vRNAs exists in the context of viral nucleoproteins and a viral RNA polymerase, forming vRNPs. Upon infection, the IAV RNA polymerase in the vRNP first transcribes the genome segments, generating viral mRNAs (**Fig. 1A-B**). This process starts with cap-snatching, which produces 9-14 nt-long capped primer from host (pre)mRNAs. The capped primers are annealed to the 3’ end of the vRNA template and extended from bases C2 or G3, depending on the sequence of the primer (18, 19). Realignment can take place on segments 4-8 when base-pairing between the capped primer and the 3’ end of the vRNA template is inefficient (19, 20). When the RNA polymerase reaches a stretch of 5-6 U-residues (U-tract) that is located 16-17 nt from the 5’ end of the vRNA template, stuttering occurs and a 3’ poly(A) tail is added to the nascent viral mRNA (21, 22) (**Fig. 1C**). The position and length of the U-tract vary slightly among the 8 IAV genome segments (**Fig. S1**), but it is not known if this impacts polyadenylation. IAV mRNAs recruit host factors UAP56, NXF1 and the TREX-2 complex to facilitate IAV mRNA export to the cytoplasm (23–25).

**Figure 1.**
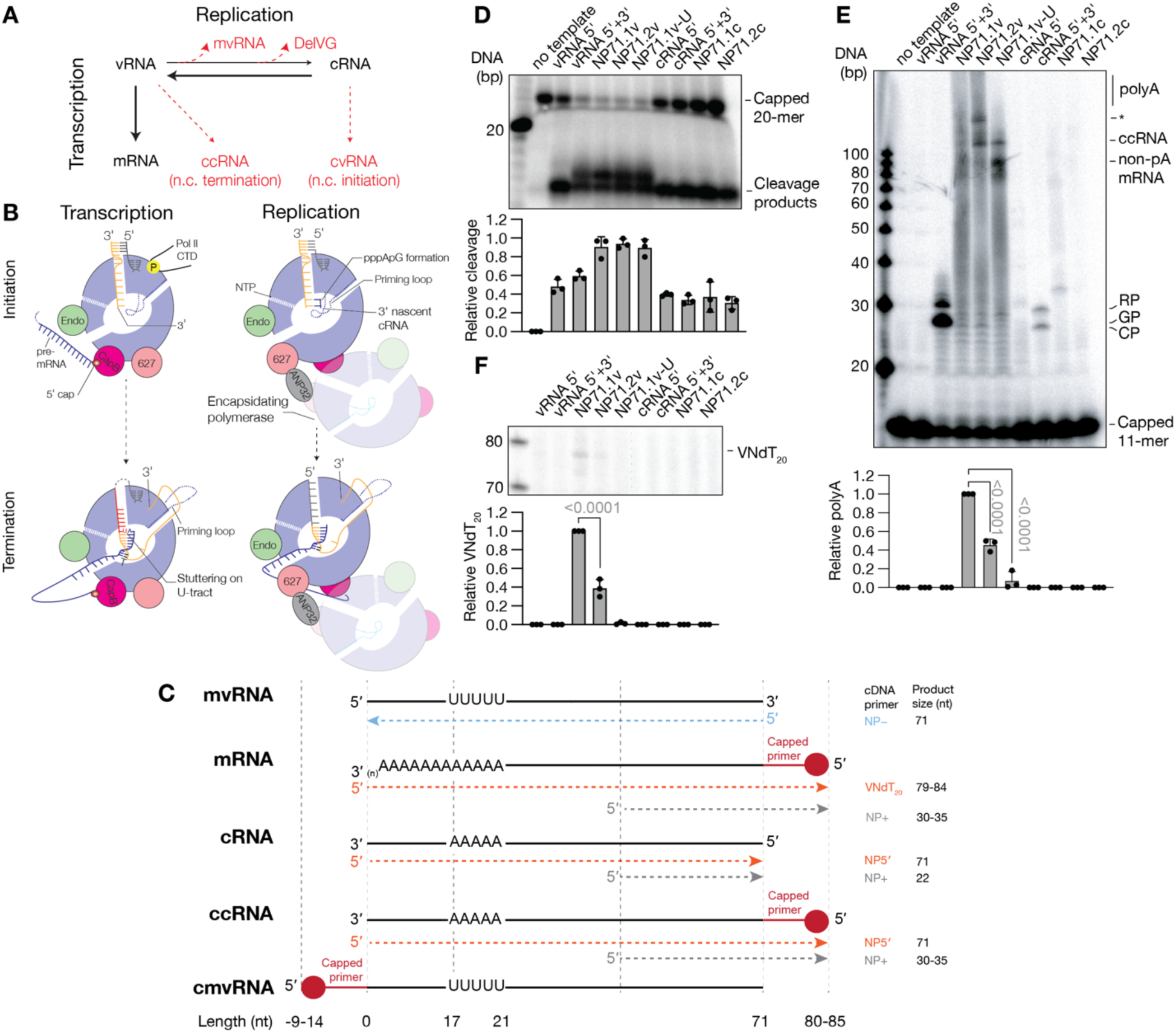
Non-canonical transcription of mvRNAs by the IAV RNA polymerase in vitro. **A**) Schematic showing the relation between canonical and non-canonical transcription and replication during IAV infection. **B**) Molecular differences between IAV transcription and replication. **C**) Overview of RNA molecules produced from mvRNA templates, and the primers used to differentiate among them. **D**) Cap-snatching of a ^32^P-labeled capped, 20-nt long RNA primer by the IAV RNA polymerase. Cleavage reactions were analyzed by denaturing PAGE. **E**) Representative image of the extension of a ^32^P-labeled capped, 11-nt long RNA primer by the IAV RNA polymerase. Extension reactions were analyzed by denaturing PAGE. The alternative initiation products CP and GP, as well as the realignment product RP are indicated. Asterisk indicates an unknown signal. **F**) Representative image of a primer extension analysis of RNA extracted from capped, 11-nt long RNA extension reactions using VNdT_20_.

In contrast to transcription, IAV replication uses *de novo* initiation mechanisms and requires host factor ANP32 (26, 27). IAV replication first produces a complementary RNA (cRNA), which is a full-length copy of the vRNA template (26–28) (**Fig. 1C**). An encapsidating RNA polymerase binds the replicating RNA polymerase to associate NP molecules with emerging an cRNA (**Fig. 1B**), creating a cRNP that subsequently serves as template for the synthesis of new vRNA molecules. cRNA and mRNA molecules thus vary in length and the sequence at their terminal ends as well as the protein context in which they exist (**Fig. 1A**). In addition to full-length molecules, IAV replication produces three non-canonical RNAs: deletion containing viral genomes (DelVG), mini viral RNAs (mvRNA), and small viral RNAs (svRNA) (29, 30) (**Fig. 1A**). DelVGs and mvRNAs lack internal genome sequences, but retain the conserved 5’ and 3’ termini of the full-length vRNA segments (29, 30). Only mvRNAs are short enough (<125 nt) to be replicated and transcribed outside the context of a vRNP or cRNP (4, 31, 32), which may facilitate their detection by RIG-I and explain the correlation between their expression and the upregulation of disease markers during infections with highly-pathogenic IAV strains. svRNA are 22-27 nt long and only contain the 5’ terminus.

mvRNA activation of innate immune signaling is not dependent on mvRNA abundance. Instead, mvRNA binding to RIG-I depends on a reduction in RNA polymerase processivity due to the formation of a template loop (t-loop) (12). However, how mvRNAs with t-loops trigger RIG-I activation remains unclear, as RNA polymerase dissociation is not affected by t-loops in cis *in vitro* (12), which precludes RIG-I binding. We here explored if non-canonical or aberrant transcription of mvRNA molecules contributes to RIG-I activation. Non-canonical IAV transcripts are capped but not polyadenylated (33, 34) (**Fig. 1A**) and referred to as capped cRNAs (ccRNA). ccRNAs can hybridize to a complementary negative sense RNA and activate RIG-I (33). We here find that mvRNAs that activate RIG-I support ccRNAs formation and that ccRNAs can trigger release of t-loop-containing mvRNAs from the RNA polymerase in a sequence-dependent manner *in vitro*. Based on our findings we propose a two-step mechanism in which sequential rounds of non-canonical replication and transcription are the start of innate immune activation by IAV RNA synthesis.

## Results

### Transcription initiation occurs on positive and negative sense mvRNAs in vitro

We previously demonstrated that the activation of RIG-I by mvRNAs was anti-correlated with the ability of the IAV RNA polymerase to efficiently replicate an mvRNA (12). To investigate whether non-canonical transcription initiation might contribute to the detection of mvRNAs by RIG-I, we explored non-canonical cap-snatching on model vRNA and cRNA promoters, as well as 71-nt long mvRNAs derived from segments 5. While non-canonical cap-snatching on cRNA promoters has been reported (35, 36), it has not been studied on mvRNA-like templates. We will refer to the product of this process as a capped mvRNA (cmvRNA; the full-length equivalent would be capped vRNA or cvRNA) (**Fig. 1A, C**).

To analyze the IAV cap-snatching activity in the presence of mvRNA templates, we purified the A/WSN/33 (H1N1) RNA polymerase (abbreviated as WSN) and incubated it with a radiolabeled, capped 20-nt long RNA in the presence of model vRNA and cRNA promoters or mvRNAs in their positive or negative sense. Specifically, we compared segment 5-derived mvRNA templates that do not (NP71.1) or do activate RIG-I (NP71.2) (12). As shown in **Fig. 1D**, the endonuclease activity was minimal when no template was added to the reaction and most efficient when the vRNA promoter or a vRNA-sense mvRNA was present. We also observed cap cleavage activity in the presence of the cRNA promoter and the positive-sense mvRNA templates (**Fig. 1D**). This non-canonical cap-snatching activity on positive-sense mvRNA templates was reproducible on mvRNAs derived from segments 2, 3, 5, 6 and 8 using either a WSN RNA polymerase or a A/Vietnam/1203/2004 (H5N1) RNA polymerase (abbreviated as VN04) (**Fig. S2**). However, the non-canonical cap-snatching activity was not dependent on the mvRNA sequence (**Fig. 1D, S2**), suggesting that this activity likely does not contribute to the differential RIG-I activation by mvRNAs.

To confirm that non-canonical cap-snatching was dependent on promoter recognition and not the rest of the mvRNA sequence, we generated alanine substitutions of PB1 C-terminal residues 699-676. Analysis of the IAV RNA polymerase transcription initiation structure showed that these PB1 residues are located on an alpha-helix near the promoter binding site and involved in interactions with base 9G of the vRNA 3’ end or U10 of the cRNA 3’ end (**Fig. S3A**), i.e., the first unpaired base that is different between the vRNA and cRNA promoter. Moreover, several previous observations suggest that mutation of PB1 699-676 could differentially impact vRNA or cRNA promoter recognition, and thus non-canonical cap-snatching: i) the PB1 helix undergoes a conformational change upon promoter binding, ii) mutation of some of the residues in the helix was shown to suppress IAV canonical cap-snatching and mRNA synthesis (37), and iii) deletion of 10A of the vRNA 5’ end, which places U10 of the vRNA 3’ end in the position of 9G (**Fig. S3A**), led to a reduction in mRNA synthesis (38). Following mutation of PB1 and purification of the mutant RNA polymerases, we observed an increase in non-canonical cap-snatching relative to wildtype (**Fig. S3B**). These results thus indicate that interactions with the promoter, rather than interactions with the downstream sequence, are important for the activation of cap-snatching and/or the dynamics of the flexible domains of the RNA polymerase (36).

### Transcription termination of mvRNAs is sequence-dependent in vitro

We next investigated whether extension of a radiolabeled, capped 11-mer primer on mvRNA templates led to non-canonical transcription termination and the production of a ccRNA. As a marker for non-canonical termination, we used 71-nt long mvRNA template in which the 5’ U-tract was interrupted with G-bases to prevent polyadenylation (NP71.1-U; **Table S1**). In all reactions, we included baloxavir (BAX) to prevent cleavage of the extended capped primers by the IAV RNA polymerase (**Fig. S4**).

On the model vRNA promoter, the RNA polymerase efficiently extended the 3’ AG of the capped primer from the 3’ C2 or 3’ G3 of the template with or without realignment at 3’ U4 (20) (**Fig. 1E, S2**). Extension of the capped primer also occurred on the model cRNA promoter by the WSN and the VN04 IAV RNA polymerase (**Fig. 1E, S2**). This non-canonical transcription initiation activity resulted in multiple products, likely because the 3’ AG of the capped primer can be aligned in two ways opposite the 3’ UC of the cRNA template and realignment may occur on a downstream U, similar to realignment on the vRNA promoter. Transcription of the NP71.1-U template generated two main termination products, one migrating at the position of a ccRNA and the other at a position of an mRNA lacking the 3’ end entirely (**Fig. 1E**), in line with our previous analyses of a segment 6 RNA with the same mutations (33). Analysis of reactions containing the negative-sense NP71.2 mvRNA template revealed signal accumulation at the ccRNA position and an interrupted polyadenylation signal (**Fig. 1E**), whereas transcription of the NP71.1 mvRNA led to a smear of polyadenylated mRNAs. These results thus suggested that a sequence in the NP71.2 template stimulated ccRNA formation (**Fig. 1E**). On the positive-sense mvRNA templates, we observed a weak extension of the capped RNA primer on all mvRNA templates tested but no difference between the NP71.1 and NP71.2 sequences (**Fig. 1E, S2**). Analysis of the RNA polymerases containing PB1 mutations revealed a differential effect on capped primer extension in the presence of a negative- or positive sense mvRNA template, in particular for PB1 N671A (**Fig. S3C**).

To confirm the presence or absence of polyA-tails in the in vitro transcription reactions, we performed primer extension reactions with a 3’ clamped oligo-dT primer (VN(dT)_20_) as described previously (33). Analysis of the reaction products showed a signal in the NP71.1 reactions and a reduced signal in the NP71.2 reactions (**Fig. 1F**). No signal was observed in the presence of any of the other mvRNA templates. Overall, these results suggest that non-canonical transcription initiation and termination can occur during IAV transcription of mvRNAs in vitro, and that these activities lead to the formation of ccRNA and cmvRNA molecules. The results further imply that only ccRNA formation is sequence dependent, while mcvRNA formation is not, and that an mvRNA that was previously shown to activate RIG-I (i.e., NP71.2) can trigger the formation of ccRNAs *in vitro*.

### mvRNAs that activate RIG-I induce ccRNA formation in cell culture

To investigate non-canonical transcription on mvRNAs in cell culture, HEK293T cells were transfected with plasmids expressing the IAV RNA polymerase subunits, NP, and mvRNA templates NP71.1, NP71.11, NP71.2, or an empty plasmid control. After twenty-four hours, we extracted RNA and used various product-specific primers to analyze the steady state RNA levels of the IAV RNA species present (**Fig. 1C**). In line with previous results (12), mvRNA templates NP71.11 and NP71.2 triggered IFN-β promoter activity in cell culture, while mvRNA template NP71.1 did not (**Fig. 2A**). Analysis of the total RNA using the vRNA-sense primer (NP-; **Fig. 1C**) showed that the NP71.11 and NP71.2 mvRNA levels were reduced and that the production of shorter aberrant products was increased (**Fig. 2B**), in agreement with our previous data (12). We did not see reproducible evidence for cmvRNA production from mvRNA templates in cell culture, even when we tried to increase cmvRNA production using the PB1 mutants (**Fig. S5**). While these results do not rule out that cmvRNA or cvRNA production may occur on other IAV RNAs in cell culture, they do further indicate that cmvRNAs are not a factor in mvRNA-dependent innate immune activation.

**Figure 2.**
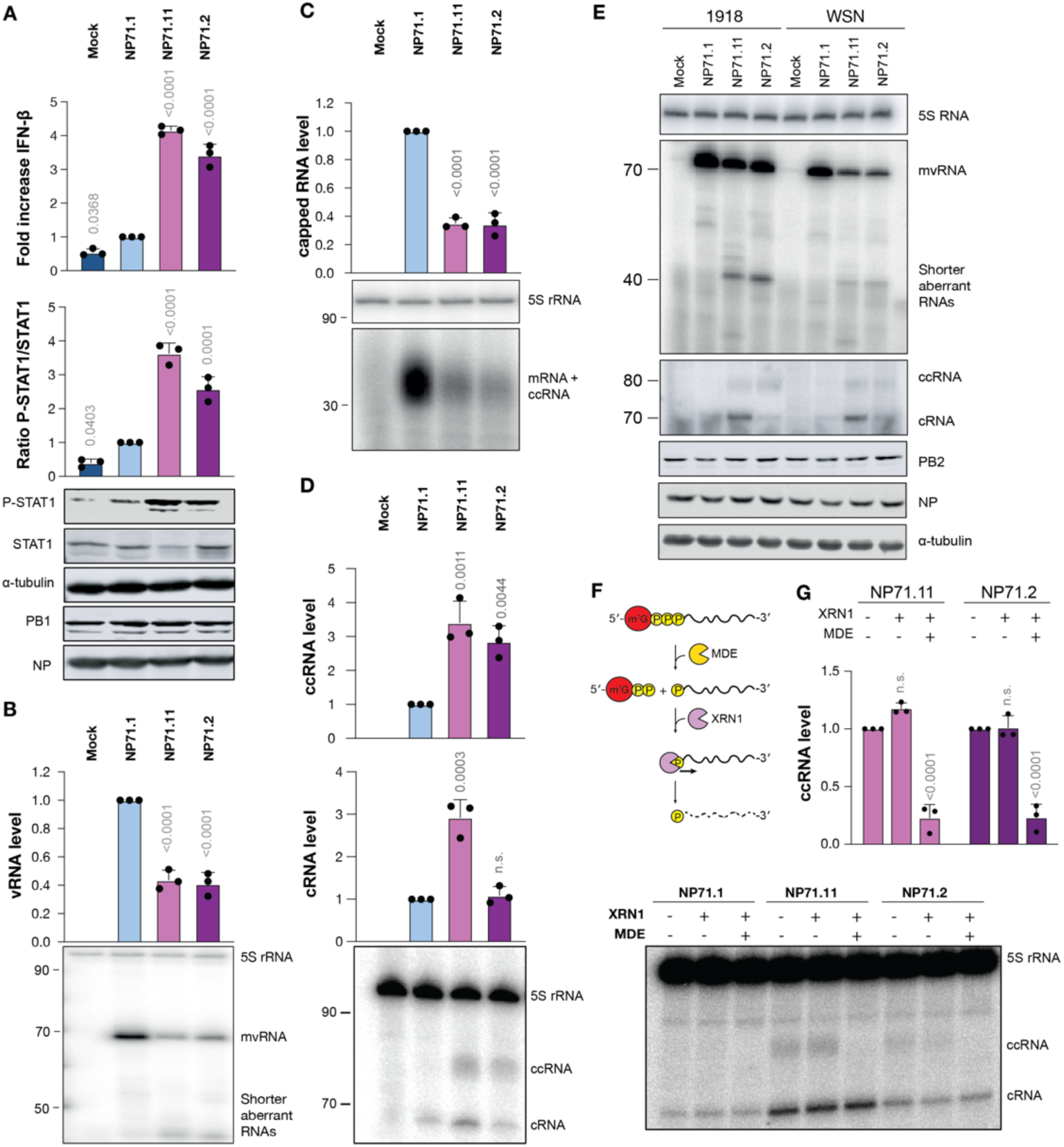
Non-canonical transcription of mvRNAs by the IAV RNA polymerase in cell culture. **A**) Innate immune activation by IAV mvRNAs in HEK 293T cells. Cells were transfected with plasmids encoding the RNA polymerase subunits, NP, a *Renilla* luciferase transfection control, a firefly-based IFN-β reporter, and the NP71.1, NP71.11 or NP71.2 mvRNAs. 24 hours post-transfection, the luciferase and phosphorylated STAT1 (pSTAT1) levels was measured. PB1, NP and tubulin protein expression was analyzed as control. **B**) Steady state mvRNA and 5S rRNA levels measured 24 hours post-transfection by primer extension using the NP-primer. **C**) Steady state total IAV capped RNA levels measured using the NP+ primer. **D**) Steady state cRNA and ccRNA levels as measured using the NP5’ primer. **E**) Steady state cRNA and ccRNA levels in the presence of the WSN or A/Brevig Mission/1/1918 (H1N1) IAV RNA polymerase. **F**) Schematic showing removal of m^7^G from the 5′ end of viral capped RNAs using mRNA decapping enzyme (MDE), generating an RNA with a 5′ end monophosphate and the release of 7-methylguanosine disphosphate. The 5′ monosphosphate RNA is then cleaved by the exoribonuclease XRN1 in a 5’ → 3’ direction, generating ribonucleotide monophosphates. **G**) Four µg of HEK 293T cell RNA was subjected to enzymatic digestion using either MDE alone or a combination of MDE and XRN1. Viral RNA levels were detected using primer extension and quantified. In all figures, graphs show mean of 3 independent experiments. Error bars indicate standard deviation and p-values were determined using one-way ANOVA.

Analysis of the total capped positive-sense RNA levels using a positive-sense-specific primer (NP+; **Fig. 1C**) yielded a smear of products due to the 9-14 nt capped primer at the 5’ end of IAV transcripts (**Fig. 2C**). Quantification of this signal showed that the steady state capped RNA level was reduced by 50% for the NP71.2 and NP71.11 mvRNAs compared to NP71.1 (**Fig. 2C**). The latter result suggests that while transcription initiation between RIG-I-activating and non-activating mvRNAs is not different in vitro, the capped RNA steady state level is significantly different between these mvRNAs in cells.

We next analyzed the steady-state cRNA and ccRNA levels using a positive-sense primer that was specific for the 3’ end of the cRNA molecules (NP5’; **Fig. 1C**). As shown in **Fig. 2D**, we observed two bands in cells expressing mvRNAs NP71.2 and NP71.11. The slower migrating of the two bands was more diffuse compared to the canonical cRNA signal and absent in NP71.1 expressing cells, suggesting that the slower migrating signal was representative of ccRNA molecules. To confirm that this signal could also be produced by an IAV RNA polymerase other than the WSN RNA polymerase, we isolated RNA produced by the A/Brevig Mission/1/1918 (H1N1) pandemic RNA polymerase (abbreviated BM18) in transfection experiments. Analysis of the steady state IAV RNA level demonstrated that the two products were also generated by the BM18 RNA polymerase on the NP71.2 and NP71.11 mvRNA templates (**Fig. 2E**).

To confirm that the slower migrating band was a transcription product and contained a 5’ cap, RNA from WSN RNA polymerase-expressing cells was subjected to an enzymatic digestion using an mRNA decapping enzyme (MDE) or a combination of MDE and exoribonuclease 1 (Xrn1) (**Fig. 2F**). MDE catalyzes the removal of 5’ cap-0 or cap-1 structures, resulting in the generation of an RNA with a 5’ monophosphate. This 5’ monophosphorylated RNA can be digested by Xrn1, whereas capped RNAs and 5’ triphosphorylated RNAs, such as cRNA and 5S rRNA molecules, are protected from Xrn1 digestion. As shown in **Fig. 2G**, in the absence of enzyme treatment or the presence of Xrn1 alone, the NP71.11 and NP71.2 reactions contained a higher molecular weight product in addition to the cRNA signal. However, in reactions that contained both MDE and Xrn1, a reduction in the higher molecular weight product was observed (**Fig. 2G**), while the cRNA and 5S rRNA loading control signals remained unaffected. These results, combined with the fact that the NP5’ primer was specific for the cRNA 3’ terminus, indicate that transcription of the NP71.11 and NP71.2 mvRNAs produces ccRNAs and that ccRNA synthesis is dependent on the mvRNA sequence in cell culture.

### RIG-I binds mvRNAs with t-loops likely in a helicase-dependent manner

In cells transfected with NP71.2 and NP71.11, activation of IFN-β promoter activity was RIG-I dependent (**Fig. S6A**), in line with previous observations (12). To test the ability of RIG-I to bind different mvRNAs, we transfected HEK293T cells with plasmids expressing the IAV RNA polymerase subunits, NP, and mvRNA templates NP71.1, NP71.11, or NP71.2. In addition, wildtype myc-tagged RIG-I (myc-RIG-I), a myc-RIG-I RNA binding mutant (K851A, K858A, K861A), or a myc-RIG-I helicase mutant (R270A) were expressed. No IFN-β promoter activity was observed in cells expressing the mutant myc-RIG-I constructs (**Fig. S6B**). After 24 hours, the steady-state RNA input and RIG-I-bound RNA levels were analyzed by primer extension (**Fig. 3A, S6C**). As shown in Fig. 3A and B, in conditions in which mvRNAs NP71.11 and NP71.2 were expressed, a 1.5-2-fold enrichment in RIG-I was found compared to NP71.1. The differential binding of the mvRNAs to RIG-I was reduced when immunoprecipitations were performed with the RIG-I RNA binding mutant and the RIG-I helicase mutant (**Fig. 3A, B**). Overall, these experiments confirm that mvRNAs that activate IFN-β promoter activity are bound more efficiently to RIG-I than an mvRNA that does not activate IFN-β promoter activity. In addition, these experiments suggest that the binding of mvRNAs NP71.11 and NP71.2 to RIG-I may involve the RIG-I helicase function, which is known to play a role in RIG-I activation and template specificity (39).

**Figure 3.**
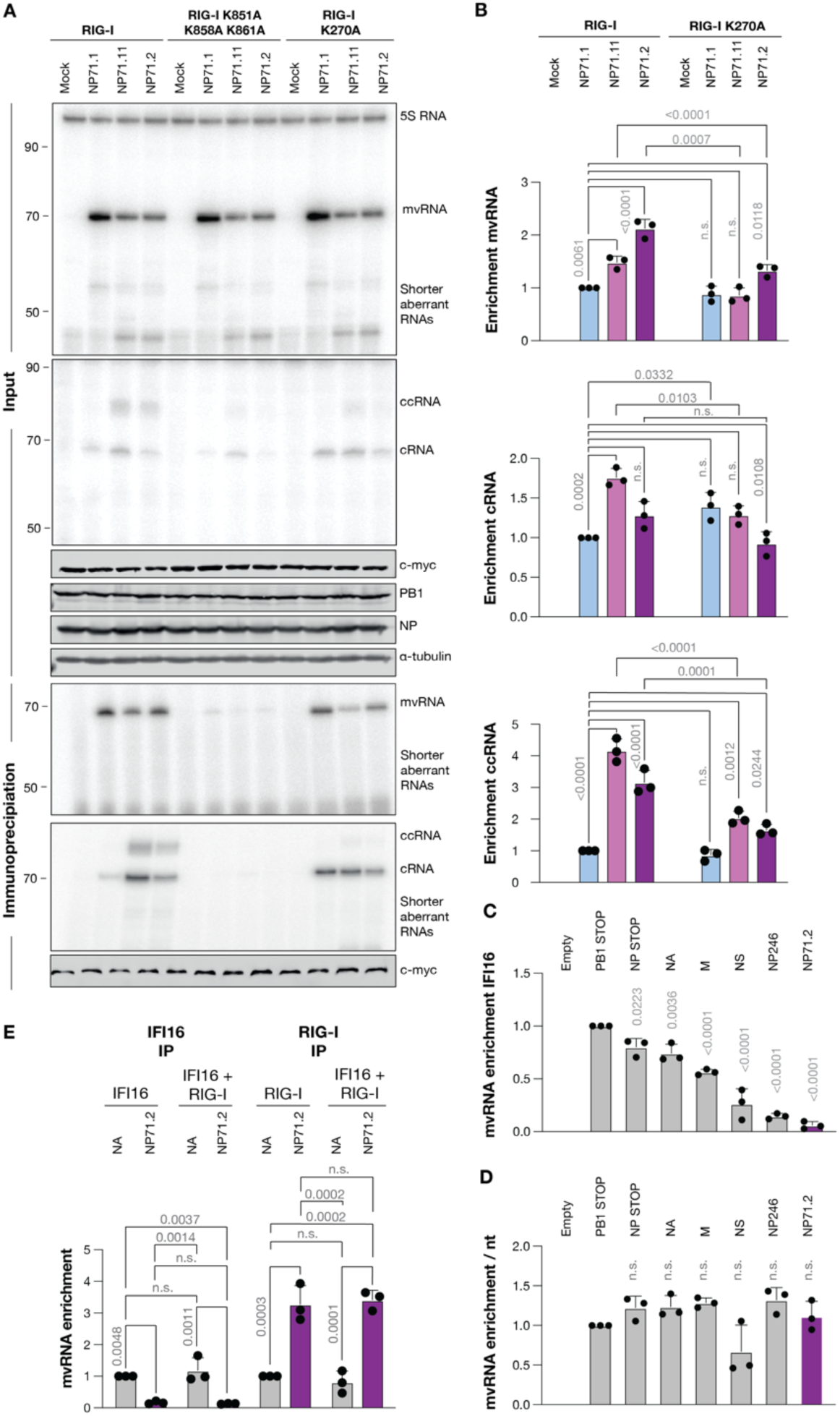
ccRNAs are enriched in RIG-I immunoprecipitation assays. **A**) RIG-I immunoprecipitation following the expression of different mvRNAs in the presence of the WSN RNA polymerase and NP in HEK 293T cells. Viral RNA levels and the 5S rRNA loading control were analyzed 24 hours post-transfection by primer extension. PB1, c-myc-RIG-I, NP, and tubulin expression were analyzed by western blot. RNA levels following immunoprecipitation are shown in the bottom panels. **B**) Quantification of the mvRNA, cRNA, and ccRNA levels following RIG-I immunoprecipitation. **C**) Quantification of the mvRNA levels following IFI-16 immunoprecipitation. **D**) Normalization of vRNA enrichment by vRNA length following IFI-16 immunoprecipitation. **E**) Quantification of the mvRNA levels following RIG-I or IFI-16 immunoprecipitation. In figures B-E, graphs show mean of 3 independent experiments. Error bars indicate standard deviation and p-values were determined using one-way ANOVA.

To investigate whether the binding of NP71.2 to RIG-I is dependent on the t-loop present in the mvRNA, we also conducted RIG-I immunoprecipitations using mvRNA NP71.6, which contains two mutations that destabilize the t-loop present in the first half of NP71.2 (**Fig. S7A**). When NP71.6 was expressed, a reduction in IFN-β promoter activity and an increase in the steady state mvRNA level was observed relative to NP71.2 (**Fig. S7B-D**), confirming previous results (12). Analysis of the RIG-I-bound RNA showed that the NP71.2 level was enriched two-fold relative to NP71.6 and NP71.1 after immunoprecipitation (**Fig. S7C, E**). Together, these data imply that destabilization of the t-loop not only influences the activity of the IAV RNA polymerase, but also RIG-I binding, in agreement with the previously observed anti-correlation between the mvRNA replication and IFN-β signaling (12). Moreover, our findings indicate that ccRNA synthesis alone is not sufficient for the activation of innate immune signaling. We will explore the latter further below.

### mvRNA-derived ccRNAs bind RIG-I

To verify whether mvRNA-derived ccRNAs bind to RIG-I, we performed immunoprecipitations using myc-RIG-I. Analysis of the input RNA levels showed clear cRNA and ccRNA signals in reactions containing the NP71.11 and NP71.2 mvRNAs, but not the NP71.1 mvRNA (**Fig. 3A; S6C**). The steady-state cRNA levels were unchanged in the RIG-I mutant reactions compared to the wildtype, except for NP71.1 in the RIG-I R270A mutant condition. In addition, we noted that the steady state ccRNA levels were consistently lower in the RIG-I binding mutant conditions compared to wildtype. This observation suggests that RIG-I binding affects either the formation or stability of these IAV RNAs in cell culture. After normalization of the RNA signals to the 5S rRNA loading control, we compared the input RNA levels to the RIG-I-bound RNA levels and calculated the enrichment relative to NP71.1. In conditions in which mvRNAs NP71.11 and NP71.2 were expressed, we observed a ∼1.5-fold enrichment in cRNA binding to RIG-I compared to NP71.1 for NP71.11, but not for NP71.2 (**Fig 3A, B**). The ccRNA signal produced by NP71.11 and NP71.2 was enriched 3 to 4-fold relative to NP71.1, although this difference was likely inflated by the reduced background in the NP71.1 immunoprecipitated signal. Both the cRNA and ccRNA signal were absent in immunoprecipitations with the RIG-I RNA binding mutant. In the immunoprecipitations with the RIG-I helicase mutant, the ccRNA signal was greatly reduced, while the cRNA signal was not. Together these results suggest that there is a correlation between the binding of innate immune activating mvRNAs to RIG-I and the binding of RIG-I to ccRNA molecules.

### IFI16 preferentially binds full-length IAV vRNAs over mvRNAs

Prior studies have shown that full-length IAV segments can bind cellular pathogen receptor IFI16 and that these interactions enhance binding of IAV RNAs to RIG-I (9). To exclude that IFI16 had affected our RIG-I-mvRNA binding observations, we transfected HEK293T cells with plasmids expressing flag-tagged IFI16 or a flag-tagged IFI16 RNA binding mutant (ΔHinA) in addition to the IAV RNA polymerase subunits and NP. As template for the RNA polymerase and RIG-I or IFI16, we co-transfected either an empty plasmid, a segment 6 vRNA (NA), or the segment 5-derived mvRNA NP71.2 (**Fig. S8**). After 24 h, the input and immunoprecipitated RNA levels were measured by primer extension. We found that the input steady-state mvRNA levels were similar between the wildtype and ΔHinA IFI16 conditions (**Fig. S8**). Analysis of the immunoprecipitated RNA showed that the ΔHinA IFI16 mutant had not bound any RNA, whereas the wildtype IFI16 had bound the segment 6 vRNA only (**Fig. S8**). Since none of the mvRNAs appear to be bound by IFI16, while the full-length NA RNA is, this suggests that IFI16 may be sensitive to nucleic acid length, as proposed for the binding of DNA by IFI16 (40).

We next explored whether RNA binding by IFI16 was correlated with the length of the IAV RNA. To this end, we expressed four IAV vRNA segments of different lengths, a segment 5-derived DelVG of 246 nt (NP246) or mvRNA NP71.2 together with flag-tagged IFI16. Following immunoprecipitation, we found that vRNA segments 2 (PB1-stop; 2341 nt) and 6 (NA; 1413 nt) were strongly enriched compared to vRNA segments 7 (M; 1027 nt) and 8 (NS; 890 nt) (**Fig. 3C, S9**). Moreover, despite the high stead-state levels of DelVG NP246 compared to the full-length vRNA segments, NP246 was not efficiently bound by IFI16. When we normalized the IFI16 binding enrichment by the RNA length, the observed differences disappeared (**Fig. 3D**), suggesting that IFI16 does not have an inherent preference for binding one IAV RNA over another, but that IFI16’s RNA binding is correlated with the number of nucleotides per RNA.

Since IFI16 was reported to enhance the interaction of a full-length vRNA segment with RIG-I, we wanted to rule out whether the presence of IFI16 could augment or suppress the binding of mvRNAs to RIG-I. To investigate this, HEK293T cells were transfected with plasmids expressing IFI16-flag, myc-RIG-I, or both in addition to the RNA polymerase subunits, NP, and either the segment 6 vRNA, mvRNA NP71.2, or an empty plasmid (**Fig. S10**). Next, immunoprecipitation of IFI16, RIG-I, or both were performed (**Fig. S10**). Analysis of the input RNA showed that the steady-state RNA levels were comparable among the different conditions (**Fig. S10**). Analysis of the IFI16-immunoprecipitated RNA showed that only segment 6 interacted substantially with IFI16, consistent with the results above (**Fig. 3E**). In RIG-I immunoprecipitations, both segment 6 and NP71.2 were found to interact with RIG-I, but NP71.2 exhibited an approximately three-fold enrichment in RIG-I binding compared to segment 6, in line previous findings (4) (**Fig. 3E**). When RIG-I and IFI16 were co-expressed, no altered NP71.2 or segment 6 binding to RIG-I was observed, suggesting that IFI16 does not stimulate or impair mvRNA binding to RIG-I (**Fig. 3E, S10**).

### ccRNA formation is correlated with U-tract truncation and not sufficient for interferon-β promoter activation

Naturally occurring mvRNAs are generated through template-switching, leading to the formation of internal deletions (**Fig. 4A**). We wondered if these internal deletions could remove residues from the U-tract. Since the U-tract is required for RNA polymerase stuttering during polyadenylation, as shown in Fig. 1 with the NP71.1-U template and by others (21, 22), a shortened U-tract could provide the simplest sequence-dependent mechanism for ccRNA synthesis. Analysis of mvRNA sequences identified in A549 cells or ferret lungs infected with three different IAV strains showed that shortening of the U-tract (i.e., 1-4 consecutive Us) occurs in ∼20% of mvRNAs (**Fig. 4A, S11A**). It is tempting to speculate that during the synthesis of some mvRNAs, the realignment step involves the 5’ U-tract if mvRNA synthesis occurs on a negative sense template (**Fig. 4A**). Alternatively, if mvRNA synthesis occurs on a positive-sense RNA template, the 3’ A-track of the cRNA may be involved. In both scenarios, realignment into the repetitive sequence would result in the truncation of the U-tract.

**Figure 4.**
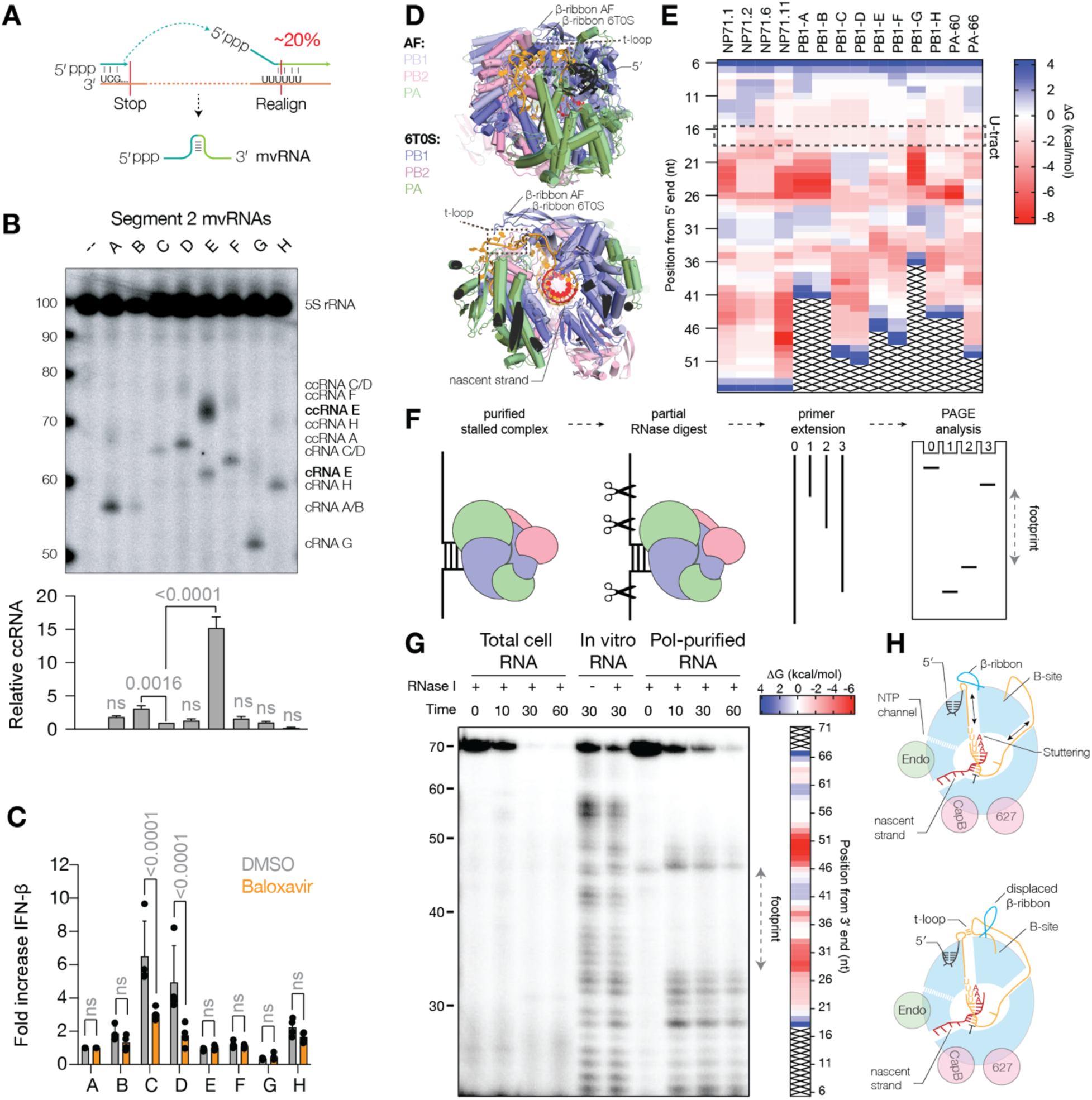
Transient RNA structures upregulate ccRNA formation. **A**) Schematic of putative mechanism for mvRNA generation using template-switching between the 3’ end of the vRNA template and the 5’ U-stretch. **B**) Primer extension analysis showing ccRNA, cRNA, and mvRNA levels in HEK293T cells expressing segment 2 mvRNAs. Graphs shows quantification of ccRNA level of three biological repeats. Error bars indicate standard deviation. P-values were calculated using One-way ANOVA relative to PB1-C. **C**) Graph of IFN-β promoter activity measuring in the presence of DMSO or baloxavir. Error bars indicate standard deviation. P-values were calculated using Two-way ANOVA with multiple corrections. **D**) Structural alignment of the IAV RNA polymerase in a polyadenylation state (PDB 6T0S) with an AlphaFold 3 (AF) model in which base pairing between the entering and exiting RNA shifts the position of the PB1 β-ribbon. **E**) Heatmap showing the stability of the t-loops in the mvRNAs analyzed. The position of the t-loop is aligned relative to the 5’ terminal end of each mvRNA to better indicate position 17 of U-tract on which polyadenylation occurs. **F**) Schematic of RNA polymerase foot printing assay. In this assay, we expressed a tandem-affinity purified (TAP)-tagged IAV RNA polymerase and an mvRNA in HEK293T cells, co-purified the mvRNA with the RNA polymerase, partially digested the mvRNA parts that are not protected by the RNA polymerase, and finally mapped the protected footprint of the RNA polymerase using primer extension. **G**) Representative image of an RNA polymerase foot printing assay in the presence of mvRNA NP71.1. Heatmap shows calculated t-loop stability for each position of the NP71.1 mvRNA. **H**) Model showing the role of the PB1 β-ribbon in polyadenylation and displacement of the β-ribbon by a transient RNA structure.

To confirm that ccRNA formation occurs on naturally occurring mvRNAs we next studied six mvRNAs derived from segment 2, PB1-A to PB1-G (12). Alignment of the segment 2 mvRNA sequences showed that mvRNA PB1-E contained a shortened U-tract (**Fig. S11B**). Analysis of the positive-sense RNA levels showed varying ccRNA signals and no correlation with IFN-β promoter activity (**Fig. 4B**). In particular, mvRNA PB1-E showed significantly increased ccRNA levels compared to the other segment 2 mvRNAs (**Fig. 4B**), indicating that U-tract shortening contributes to ccRNA formation. The mvRNA levels varied in accordance with the same t-loop stability described previously (**Fig. S11C**) (12) and the most suppressed mvRNAs (C and D) showed the highest IFN-β promoter activity (**Fig. 4C**). To test if transcription contributed to IFN-β promoter induction by mvRNA C and D, we treated transfection reactions with PA endonuclease inhibitor BAX and observed a significant reduction in IFN-β promoter activity (**Fig. 4C**).

To confirm that transcription plays a role in innate immune activation by mvRNAs during infection, we overexpressed the viral RNA polymerase during WSN infection. As shown previously, this creates an imbalance between viral RNA polymerase and NP levels, and stimulates mvRNA synthesis and IFN promoter activation (4) (**Fig. S11D**). Addition of BAX 2 h post infection to conditions in which mvRNAs were expressed reduced IFN promoter activation without significantly reducing mvRNA steady state levels (**Fig. S11D**). We note that these infection experiments can never be fully conclusive, since BAX also affects the expression of viral proteins implicated in modifying the host response to virus infection (e.g., NS1, PB1-F2, PA-X). We are therefore cautious and conclude that the infection results indicate that there is a correlation between transcription and innate immune activation, in line with observations by others (41, 42). Overall, the results indicate that ccRNA synthesis alone is not sufficient and a reduction in RNA polymerase processivity (i.e., reduced mvRNA replication) is needed as well. These observations are in line with our previous analysis showing that ccRNA molecules alone are not sufficient to activate RIG-I (33). Similarly, a reduction in RNA polymerase processivity on mvRNAs appears not to be sufficient for RIG-I activation and a transcriptional component is needed to contribute to the induction of IFN-β promoter activity.

### The formation of ccRNAs is correlated with the presence of a terminal t-loop

We and others recently demonstrated that disruption of triple-stranded β-sheet formation interrupts polyadenylation and triggers ccRNA formation (33, 34). As ccRNA formation on mvRNAs was sequence dependent, we wondered if a transient RNA structure could interrupt polyadenylation by repositioning the PB1 β-ribbon. An AlphaFold 3 (43) model using the WSN RNA polymerase and an mvRNA template with a stabilized t-loop over the 5’ U-tract implied that the PB1 β-ribbon could be displaced upwards by a t-loop (**Fig. 4D**). We next performed a sliding-window RNA folding analysis of the mvRNAs tested above as described previously (12) and presented the calculated ΔG in a heatmap (**Fig. 4E**). Our analysis of the negative-sense sequences showed that t-loop formation can potentially occur on all mvRNAs tested above, but not on NP71.1. For comparison we also included two segment 3 mvRNAs, one of which is known to activate RIG-I (PA-66) while the other one is not (PA-60). We observed that the t-loop pattern of PA-60 was similar to NP71.1, while PA-66 was similar to NP71.2 (**Fig. 4E**). Analysis of the ccRNA level showed that PA-66 triggers higher ccRNA level production than PA-60 (**Fig. S12A**) and induced higher IFN-β promoter activity (**Fig. S12B**).

We next aimed to experimentally show that t-loops can form in cell culture using an RNA polymerase foot-printing assay (**Fig. 4F**). Analysis of the RNA polymerase footprint on the NP71.1 mvRNA revealed an undigested region between 33-49 nt that was flanked by cleavage signals (**Fig. 4G**). These signals were not present when RNase I or the RNA polymerase were absent. Destabilization of the t-loop (**Fig. S12C**) using single or double mutations changed the intensity of the footprint signal (**Fig. S12D-E**). The same footprint was obtained when we used MNase to digest unprotected RNA (**Fig. S12F**). We found that the RNase protected sequences mapped closely to the predicted t-loop sites for mvRNA template NP71.1 (**Fig. 4G, S12C**), indicating that our sliding window analysis captures the stalling of the RNA polymerase *in vitro* (12) and in cell culture.

On the co-purified NP71.2 and NP71.11 mvRNAs, no clear digestion pattern was observed (**Fig. S12G-H**), preventing us from confirming t-loop formation around the U-tract. In addition, we noticed that reproducibly less RNA co-purified with the RNA polymerase, either due to reduced template amplification and/or increased template release in cell culture. While we can presently not fully rule out that the NP71.2 and NP71.11 templates contain alternative structures that could have prevented RNase digestion, the results above are overall indicative of a model in which t-loop formation near the U-tract displaces the PB1 β-ribbon and interrupts polyadenylation (**Fig. 4H**).

### mvRNA-derived ccRNAs activate RIG-I as part of a heteroduplex

We recently showed that ccRNAs do not induce RIG-I activation directly, but need to hybridize to a complimentary RNA, such as an mvRNA or small viral RNA (svRNA) (33). To confirm that this model holds true for mvRNA-derived ccRNAs, we used the PA-66 mvRNA to generate mvRNA, svRNA and ccRNA T7 transcripts. Incubation of purified RIG-I with these in RNA molecules only resulted in ATPase activity in the case of 5’ triphorphorylated mvRNA molecules, whereas the ccRNA or alkaline phosphatase-treated mvRNA molecules did not trigger ATPase activity (**Fig. S13**). However, when we combined the ccRNA molecules with a 5’ triphosphorylated svRNA, we observed a significant increase in ATPase activity compared ccRNA or svRNA molecules alone (**Fig. S13**). Note that we used the short svRNA-like molecule, since the mvRNA alone already activated RIG-I. Together, these results suggest that mvRNA-derived ccRNAs need to hybridize with a negative-sense molecule that contains a 5’ triphosphate to bind and activate RIG-I, in agreement with what we reported previously (33).

### The presence of ccRNAs can trigger mvRNA template release in vitro

To form a duplex between a ccRNA and an mvRNA, the mvRNA template must be released by the IAV RNA polymerase. We previously showed that RNA polymerase stalling is not sufficient for mvRNA release (12). To test if ccRNAs can contribute to mvRNA release in a sequence-dependent manner, we immobilized an mOrange-tagged IAV RNA polymerase on magnetic beads and added a radiolabeled mvRNA template (**Fig. 5A**). Excess template was removed through washing and the replication reaction started by adding NTPs, MgCl_2_, ApG, an encapsidating inactive RNA polymerase (PB1a) as described previously (12). A ∼2x excess of complementary ccRNA or a non-complementary ccRNA derived from NA-71 (33) was added to test template release. After a 30 min incubation, we separated the free and bound fractions through centrifugation and quantified the level of template mvRNA present in each fraction using dot blotting (**Fig. 5B**). We found that the presence of a complementary ccRNA, but not a ccRNA that lacked complementarity, significantly increased the mvRNA signal in the unbound fraction (**Fig. 5B**). Finally, we analyzed the unbound fraction non-denaturing PAGE to assess duplex formation between the radiolabeled mvRNA and the complementary ccRNA. As shown in Fig. S14, this analysis showed that the mvRNA template is able to form a higher molecular weight product in the presence of a ccRNA in the unbound fraction relative to the ccRNA in the reaction that lacked ccRNA (**Fig. S14**). Taken together, these findings indicate that, at least in vitro, ccRNAs can contribute to the release of an mvRNA template in a sequence-dependent manner (**Fig. 5A** and **C**). Unfortunately, we were not able to confirm the steps of the mechanism in cells, because expression of a negative-sense RNA template from a pPol-I plasmid in the presence or absence of RNA polymerase and NP prior to ccRNA addition is already sufficient to activate RIG-I (33).

**Figure 5.**
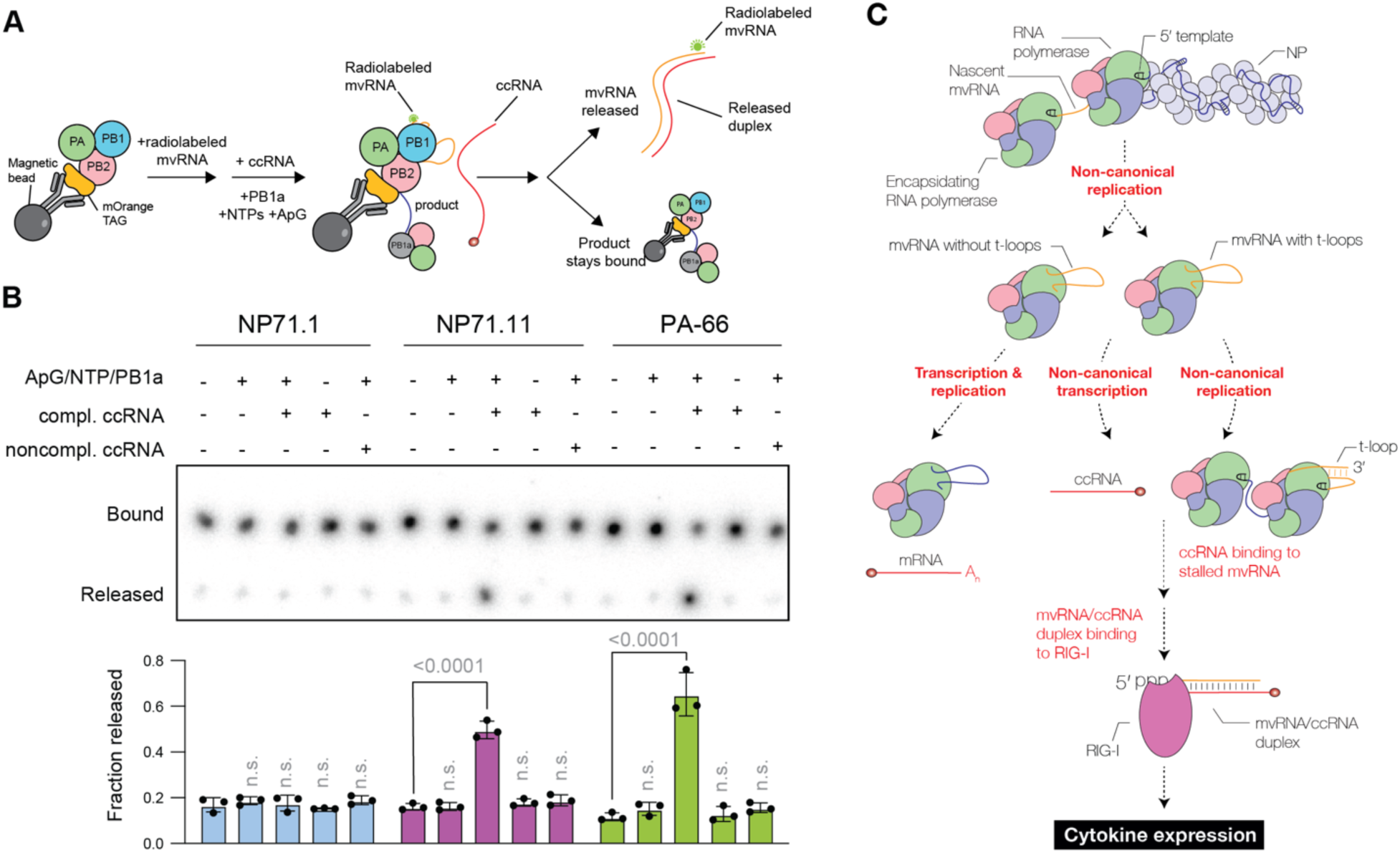
ccRNAs trigger release of mvRNA template in a sequence-dependent manner. **A**) Schematic of assay to test impact of ccRNA molecules on mvRNA template release during replication *in vitro*. Purified RNA polymerase is immobilized on magnetic beads and incubated with radiolabeled mvRNA template. Excess template is washed away and ApG, NTPs, and an inactive RNA polymerase (PB1a) are added to start replication. A complementary ccRNA or non-complementary ccRNA derived from a segment 6 mvRNA (NA71) are added to trigger template release. **B**) Dot blot of RNA polymerase-bound and unbound fractions. Graph shows signal measured using radiography. **C**) Model of RIG-I activation after non-canonical replication and transcription of t-loop-containing mvRNAs. mvRNAs that do not contain a t-loop that affects replication may contribute to RIG-I activation by mvRNAs derived from the same segment *in trans* if they support ccRNA formation.

## Discussion

RIG-I is preferentially activated by 5ʹ tri- or diphosphorylated dsRNA molecules (39), but relatively few dsRNA molecules are produced during IAV infection (7, 44). Instead, IAV RNA binding to RIG-I has been proposed to occur via the partially complementary 5’ and 3’ termini of each RNA. However, all IAV RNA segments and non-canonical RNAs have near-identical terminal ends, and innate immune activation by DelVGs as well as mvRNAs is sequence-dependent (12–14). The precise mechanism through which some IAV RNAs become more potent inducers of innate immune signaling than other IAV RNAs remains unclear (12–14, 45). Since aberrant positive sense RNA virus transcripts can contribute to the activation of IFN signaling (46, 47) and non-canonical IAV transcripts have been described, we here investigated the impact of non-canonical transcription on the differential detection of mvRNAs by RIG-I.

We observed that mvRNAs can trigger ccRNA formation in a sequence-dependent manner and propose that ccRNA formation occurs when the 5’ U-track in the mvRNA is shortened during mvRNA formation or when the mvRNA contains a t-loop that dysregulates the function of the PB1 β-ribbon. This means that t-loops are implicated in a further IAV process in addition to contributing to multi-basic cleavage site emergence in the hemagglutinin-encoding gene (48), reducing replication (12), and altering genome segment stability (49). We further found that mvRNAs that i) contain a t-loop in the first half of the RNA sequence and ii) can trigger ccRNA synthesis are enriched in RIG-I binding experiments compared to mvRNAs that lack these characteristics, even when the latter mvRNAs were more abundant in cells due to increased replication. We confirmed that mvRNA-derived ccRNAs only activate RIG-I when a complementary RNA with triphosphate is present, as we showed elsewhere (33), and in line with the binding preference of RIG-I for a 5’-ppp and a blunt dsRNA end (39). Moreover, the importance of mvRNA transcription for RIG-I activation corroborates findings that link IAV transcription to innate immune activation (41, 42).

RIG-I binding alone is a poor predictor of RIG-I activation. Prior research has demonstrated that RNA sampling and RIG-I activation involve multiple checkpoints, including RNA recognition, ATP binding, and ATP hydrolysis (50–53). In these steps, ATP binding stabilizes RIG-I complexes, while ATP hydrolysis destabilizes them. Since RIG-I is deprived of ATP during immunoprecipitation, the experimental procedure likely freezes RIG-I in a state where the dissociation of an mvRNA cannot occur. A highly abundant mvRNA, like NP71.1, can thus be immunoprecipiated with RIG-I even when this mvRNA does not activate RIG-I. However, a low abundant mvRNA that activates RIG-I will become enriched over time. Consistent with this analysis, mutation of the RIG-I helicase/ATPase domain leads to IAV RNA immunoprecipitation without enrichment for IAV RNAs that activate innate immune signaling. In this condition, only limited RIG-I dissociation occurs and the immunoprecipitation therefore reflects abundance instead of activation. Our results agree with the above selective mechanism.

We previously showed that IAV RNA polymerase stalling does not result in the release of the RNA template or the mvRNA products *in vitro*. Our foot printing results confirm that once the RNA polymerase enters a strong t-loop the RNA polymerase does indeed not dissociate from the template RNA. Interestingly, we find that the presence of a ccRNA can cause template RNA release, which may lead to duplex formation (**Fig. 5C**). This duplex may be bound by RIG-I or NS1 (54), similar to the dsRNA structures recently identified by atomic force microscopy during IAV replication *in vitro* in the absence of free NP (55).

Finally, we observed that IFI16 does not play a role in mvRNA binding or improving the binding affinity of mvRNAs to RIG-I. Instead, we showed that the binding of IFI16 to IAV RNA is largely length dependent. Previous studied have shown that IFI16 requires a minimum of 50-70 base pairs of exposed dsDNA to form filaments (40). Our results indicate that IFI16 also requires a minimal RNA length and that mvRNAs are not efficiently bound by IFI16. We therefore propose that mvRNA-ccRNA duplexes predominantly bind to RIG-I directly, without the involvement of IFI16. In contrast, full-length segments interact with both IFI16 and RIG-I, potentially engaging in competitive binding for the RNA segments, as previously suggested (56). While we did not test the impact of mvRNA-ccRNA duplex formation on IFI16 binding, we expect this not to have been a differentiating factor since IFI16 can bind dsDNA. Overall, our results indicate that different IAV RNA species interact with host innate immune receptors in different steps or different pathways, and that non-canonical transcription as well as modulation of RNA polymerase processivity during replication is required for activation of the innate immune response by mvRNAs (**Fig. 5C**).

## Methods

### Cells and transfection

HEK293T and MDCK cells were obtained from the American Type Culture Collection (ATCC) and confirmed to be free of mycoplasma. HEK293 and HEK293 RIG-I -/- cells were a gift from J. Rehwinkel (Oxford University) and have been described previously (57). All cells were grown in Dulbecco’s Modified Eagle Medium (DMEM) High Glucose with L-glutamine and pyruvate (Gendepot), supplemented with 10% fetal bovine serum (FBS HI, Gibco Life Technologies). The cells were maintained at 37°C and 5% CO_2_. Cells were counted with a Countess 3 automated cell counter (Invitrogen) prior to experiments.

### Plasmids

PB1 mutations described in this study were introduced using site-directed mutagenesis using primers listed in Table S2. The pPolI plasmids encoding A/WSN/33 genome segments were previously described in (58); the pPolI NP246 plasmid was described in (4); the pcDNA3 plasmids encoding wild-type A/WSN/33 proteins in (59); the pcDNA3 plasmid encoding PB1 D445A/D446A (PB1a) mutant in (60); the pcDNA3 plasmid encoding myc-RIG-I wildtype and mutants in (4, 33); the pcDNA3 plasmid encoding TAP-tagged PB2 in (61); the plasmid expressing firefly luciferase under IFN-β promoter (pIFD(−116)lucter) and pcDNA3 encoding *Renilla* luciferase in (12); pcDNA3 plasmids encoding flag-tagged IFI16 wildtype and mutant were described previously (62) and obtained from Addgene (plasmid numbers 35064 and 35062).

### Transfections

The RNP reconstitution assays were performed as described previously (63). Briefly, confluent cells were trypsinized, washed with PBS, and resuspended in 10 ml of DMEM/10% FBS. Cells were counted and 4×105 HEK-293T cells were seeded in a 24-well plate, and transfected in suspension with 250 ng of the WT plasmids pcDNA3-NP, pcDNA3-PA, pcDNA3-PB2, WT or mutated pcDNA3-PB1, and a pPolI plasmid encoding a vRNA full-length or truncated vRNA/cRNA templates (**Table S2**) using Lipofectamine 2000 (Invitrogen) according to the manufacturer’s instructions. Next, 500 μl of DMEM/10% FBS containing 1% Penicillin/Streptomycin (Gibco) was added to each well. The cells were incubated for 24 h, resuspended in 1 ml PBS, and split in three equal parts. Five hundred μl of cell suspension was used for RNA isolation, 250 μl for western blot analysis and another 250 μl for the IFN-β promoter expression assay. Cells were pelleted at 700 x g for 5 min and analyzed.

### IFN-β promoter

For IFN-β promoter activity assays, 100 ng/well of a plasmid expressing firefly luciferase from the IFN-β promoter and 10 ng/well of a plasmid expressing *Renilla* luciferase from the CMV promoter were transfected alongside IAV expression plasmids. Twenty-four hours after transfection, cells were collected and a quarter of each pellet resuspended in 50 μl PBS. Next, 25 μl of cell suspension was transferred in duplicate to a white 96-well plate (Greiner Bio-One) and mixed with 25 μl substrate and lysis buffer. The firefly luciferase and *Renilla* luciferase activities were measured using a Dual-Glo Luciferase Assay System (Promega) according to the manufacturer’s protocol. The signals were measured on a Synergy LX (Bio-TEK) plate reader. The signal of non-transfected cells was used as background and subtracted from firefly luciferase and *Renilla* luciferase signals. Firefly luciferase signal was divided by the *Renilla* luciferase signal to normalize the firefly luciferase signal for the transfection efficiency.

### Virus and infection

A/WSN/33 (H1N1) was rescued using the 12-plasmid system (58). Virus stocks were grown on MDCK cells at an MOI of 0.01. For BAX experiments, HEK293T cells were transfected (250 ng) with pcDNA3 plasmids expressing the A/WSN/33 (H1N1) RNA polymerase or an empty pcDNA3 plasmid, 100 ng IFN-β reporter plasmid, and 10 ng *Renilla* transfection control plasmid. After 24 h, the cells were infected or mock infected with A/WSN/33 (H1N1) at an MOI of 3. After 2 h, BAX was added to a final concentration of 30 nM in DMEM/0.5% FBS. Eight hours post infection, the cells were harvested and the firefly and *Renilla* luciferase signals measured. RNA was extracted for mvRNA RT-PCR as described previously (4).

### RIG-I activity assay

Recombinant RIG-I was purified as described in (64). For ATPase assays, 0.5 µM RIG-I was incubated with 0.1 µM [γ-32P]ATP and 10 ng of template RNA as previously described (33). [γ- ^32^P]ATP and ^32^P_i_ were resolved using PEI-cellulose TLC plates (Sigma-Aldrich). After wrapping the TLC plates in plastic foil, the radioactive signals were detected using phosphorimaging.

### RNA extraction

Cell pellets were resuspended in 250 µl TRI Reagent (MRC) and mixed in a 5:1 ratio with chloroform by vortexing. The suspension was centrifuged for 15 min at 10000 x g and 4 °C. The aqueous phase (approximately 150 μl) containing the RNA was transferred to a new tube, mixed with 1.5 μl of GlycoBlue TM Coprecipitant (Thermofisher) and 1 volume (approx. 150 μl) of isopropanol. After mixing by inversion, the RNA was centrifuged for 15 min at 17000 x g and 4 °C. The RNA pellet was subsequently washed with 75% ethanol and centrifuged for 5 min at 17000 x g and 4 °C. Residual ethanol was removed by a further 10 second centrifugation step. Finally, RNA pellets were resuspended in 10 μl of RNase-free water and the RNA concentration measured by the NanoDrop spectrophotometer (Thermo Scientific).

### Primer labeling

One μM of DNA oligonucleotide (Table S3) was incubated with 10 U T4 polynucleotide kinase (NEB), 1x buffer T4 polynucleotide kinase, and 1 μl [γ-32]P-ATP (6000Ci/mmol; Perkin Elmer) in a total reaction volume of 10 μl at 37°C for 1 h. Unincorporated radioactive label was removed using an Oligo Clean & concentrator Kit (Zymo Research) according to the manufacturer′s instructions, and radiolabeled oligonucleotides were eluted in 30 μl RNase-free water.

### Primer extension

For quantitative primer extensions, reverse transcription was carried out using SuperScript III reverse transcriptase (ThermoFisher Scientific) with ^32^P-labelled oligonucleotides (Table 3; Fig. 1) that were complementary to IAV vRNA and ribosomal 5S rRNA (5S rRNA). First, 2-4 μl of total RNA was annealed to 1 μl of the radioactive primer mix (0.25 μl of 0.3 μM ^32^P-primer, 0.05 μl of 0.3 μM ^32^P-5S_100 primer, 0.45 μl of 10 μM unlabeled 5S_100 primer, and 0.25 μl water) incubated at 95 °C for 2 min and immediately cooled on ice. Reverse transcription was conducted in a 10 μl reaction volume using 1X First-strand buffer, 50 U SuperScript III Reverse Transcriptase (Invitrogen), 10 mM dithiothreitol (DTT), 10 U of RNase inhibitor (APExBio), and 0.5 mM dNTPs by incubating RNA samples at 50 °C for 1 h. Ten μl of formamide loading dye (90% Formamide, 10 mM EDTA, 0.25% Bromophenol Blue, and 0.25% xylene cyanol FF) was added and the samples denatured at 95 °C for 2 min. The ^32^P-labeled cDNA products were resolved by 6% or 12% denaturing PAGE (19:1 acrylamide/bisacrylamide (Bio-Rad), 1x tris borate buffer (TBE, Growcells), 7 M urea, 0.1% APS, 0.1% TEMED) for 1 h at 35 W and 2000 V. The 6% denaturing PAGE gels were dried on chromatography paper grade 1 (Cytiva) using a gel dryer (Model 583, Bio-Rad) at 80°C, for 1 h. The radiolabeled signals were measured using phosphorimaging on a typhoon scanner (GE Healthcare). The 5S rRNA signal was used to normalize the viral RNA levels.

### Decapping

To assess the presence of a m^7^G cap on the extended cRNA products, 4 μg of total cellular RNA was incubated in a 40-μl reaction mix with 1X mRNA decapping enzyme reaction buffer, 15 U of mRNA decapping enzyme (NEB), 40 U of RNAse inhibitor (APExBio), and incubated at 37 °C for 2 h. RNA was purified using a RNA clean and concentrator 5 kit (Zymo Research) and eluted in 15 μl RNase free water. In a 50-μl reaction mix, 15 μl of the eluted RNA was further digested in the presence of 1X NEBuffer 3, 50 U of RNAse inhibitor (APExBio), and 5 U of XRN-1 (NEB). These reactions were incubated at 37 °C for 2 h. Finally, the RNA was purified using a RNA clean and concentrator 5 kit (Zymo Research) and eluted in 10 μl for primer extension analysis.

### Western blot

Protein samples were resuspended in 5X Laemmli buffer (For 5X; 300 mM Tris-HCl pH 6.8, 10% SDS, 50% glycerol, 0.05% bromophenol blue and 100 mM DTT) and denatured at 95°C for 5 min. Insoluble debris was pelleted through centrifugation at 17000 *x g* for 5 min. The SDS-PAGE gels consisted of two parts: a bottom 8% resolving gel (375 mM Tris-HCl pH 8.8, 0.1% SDS, 8% acrylamide/bis 37.5:1 (Bio-Rad), 0.1% APS, 0.1% TEMED) and a 3.2% top stacking gel (125 mM Tris-HCl pH 6.8, 0.1% SDS, 3.2% acrylamide/bis 37.5:1 (Bio-Rad), 0.1% APS, 0.1% TEMED). Samples were run at 100V in 1X Tris-glycine SDS running buffer (Invitrogen). Proteins were transferred to a Protan 0.45 μM (Cytiva) at 25V, 1A for 25 min using transfer buffer (25 mM Tris, 192 mM glycine, and 20% ethanol), Whatman gel blotting paper (Cytiva) and a Trans-Blot Turbo transfer system (Bio-Rad). The membranes were first incubated with blocking buffer (PBS-1X, 5% BSA, 0.1% Tween-20) for 1 h. The membranes were incubated overnight at 4°C with primary antibodies (Table S3) diluted in blocking buffer. Before adding secondary antibodies at 4°C for 2 h (Table), the membranes were washed three times with PBS 0.05% Tween-20 for 10 min. Membranes were washed again three times before the signals were measured on an Odyssey CLx (LI-COR).

### RNA polymerase purification

Twenty-four h before transfection, 5.5×10^6^ HEK 293T cells were plated in a 10 cm dish. The cells were transfected with 4 μg of pcDNA3 plasmids expressing the PB1 subunit, the PA subunit and a TAP-tagged PB2 subunit, using 1 mg/ml PEI with a 1:5 DNA:PEI (µg:µl) ratio. Forty-eight h after transfection, the cells were washed 2 times with cold PBS, lysed with 1 ml of lysis buffer (50 mM Hepes pH 8, 200 mM NaCl, 25% glycerol (Sigma #G5516-1L), 2% Tween-20 (RPI #9005-64-5), 1 mM β-mercaptoethanol (Bio-Rad), and 1X EDTA-free Protease inhibitor cocktail (Roche)) directly in the dish and harvested. The cells were further lysed by rotation for 1 h at 4°C before being sonicated. The lysate was centrifuged at 17000 x *g* for 15 min at 4°C. The TAP-tagged RNA polymerase was bound to 50 μl IgG Sepharose beads (GE Healthcare) that were pre-washed 3 times in binding buffer (50 mM Hepes pH 8, 200 mM NaCl, 25% glycerol, 2% Tween-20). After 24 h of continuous rotation at 4°C, the beads were washed 3 times in binding buffer and 1 time in cleavage buffer (50 mM Hepes pH 7.5, 200 mM NaCl, 25% glycerol, 0.5% Tween, 1 mM DTT). Each wash was followed by 10 min rotation at 4°C. Finally, Tobacco etch virus (TEV) protease cleavage was conducted with 15 U of AcTEV protease (Invitrogen, #12575-015) in 250 μl of cleavage buffer overnight. The beads were separated from the cleaved RNA polymerase by centrifugation at 500 *x g* for 1 min and the partially purified RNA polymerase was analysed through SDS-PAGE and silver staining using a SilverXpress kit (Invitrogen).

### RNA polymerase foot printing

Transfections were performed as described above for the RNA polymerase purification with minor modifications. The cells were transfected with 4 μg of pcDNA3 plasmids expressing the wildtype PB1 and PA subunits, a TAP-tagged PB2 subunit, and a pPolI plasmid expressing an mvRNA template using a 1:5 DNA:PEI (µg:µl) ratio. Forty-eight h after transfection, the cells were harvested in cold PBS, washed once with cold PBS, and lysed with 1 ml of EDTA lysis buffer (50 mM Hepes pH 8, 5 mM EDTA, 200 mM NaCl, 25% glycerol, 2% Tween-20, 1 mM β-mercaptoethanol, and 1X EDTA-free Protease inhibitor cocktail). Lysis was continued for 1 h at 4°C and completed using sonication. The lysate was cleared through centrifugation at 17000 x *g* for 15 min at 4°C. The TAP-tagged RNA polymerase bound to 50 μl IgG Sepharose beads (GE Healthcare) that were pre-washed 3 times in lysis buffer for 2 hours at 4°C under continuous rotation. To remove background RNA, the beads were washed three times with lysis buffer. Finally, RNase I (Thermo Scientific #EN0602) was added into the lysis buffer using the amounts indicated for the experiments or micrococcoal nuclease (Thermo Scientific #EN0181) was added in the manufacturer’s reaction buffer as indicated for the experiments. Undigested RNA was extracted using 250 µl TRI Reagent (MRC) as described above.

### Immunoprecipitation

Twenty-four h before transfection, 5.5×10^6^ HEK 293T cells dish were plated in a 10 cm dish. The cells were transfected with 4 μg of pcDNA plasmids expressing the PB1 subunit, the PA subunit and a PB2 subunit and 4 μg of pcDNA c-myc RIG-I or/and pcDNA c-myc IFI-16 and mvRNA or full length segments encoded by a pPol-I plasmid, using 1 mg/ml PEI (Sigma) with a ratio 1:5 DNA:PEI. 48h after transfection, the cells were washed 2 times with cold PBS, lysed with 600 μl of lysis buffer (10 mM Tris/HCl pH7.5, 200 mM NaCl, 0.5% NP40, 1X EDTA-free Protease inhibitor cocktail (Roche)) directly in the dish, harvested with a cell scraper, and incubated 1 h at 4°C on a rotation wheel before being sonicated. The lysate was centrifuged at 17000 *x g* for 15 min at 4°C. The lysates were further diluted with 900 μl of dilution buffer (10 mM Tris/HCl pH7.5, 200 mM NaCl, 1X EDTA-free protease inhibitor cocktail (Roche)). The c-myc-RIG-I or Flag-IFI16 were bound to 25 μl of Myc-TRAP agarose (Chromotek) or Fab-TRAP agarose (Chromotek) that were pre-washed 3 times in wash buffer (10 mM Tris/HCl pH7.5, 200 mM NaCl, 0.05% NP40). Each wash is followed by 10 min of rotation at 4°C. Beads were resuspended in 100 μl of wash buffer. Eighty μl of this resuspension was used for RNA extraction and 20 μl for protein analysis through SDS-PAGE and western blot analysis.

### Primer capping

The capping of RNA primers was conducted as described previously (27). Briefly, 1 μM of 20-nt-long RNA (ppAAUCUAUAAUAGCAUUAUCC; Chemgenes) or 11-nt-long RNA (ppGAAUACUCAAG) were capped with a radiolabeled cap-1 structure in a 10 μl reaction containing 0.15 μM [α-^32^P]GTP (3000 Ci mmol^−1^; Perkin-Elmer), 1X Capping buffer (NEB), 0.8 mM S-Adenosylmethionine (SAM) (NEB), 5 U vaccina virus capping system (NEB, M2080S), 25 U mRNA Cap 2’ O-methyltransferase (NEB, M0366S) for 1h at 37°C, 4 min 70°C. Unincorporated radiolabel was removed with the Oligo Clean & concentrator Kit (Zymo Research) according to the manufacturer′s instructions, and radiolabelled RNAs eluted in 30 μl of RNAse-free water.

### Cap cleavage

The cleavage of RNA primers by the IAV RdRp was conducted as described previously (27). To test the endonuclease activity of the purified IAV RdRp, in a 4 μl reaction ^32^P-labelled and capped 20-nucleotide-long RNA primer was incubated with 1 mM DTT, 0.7 μM template (Table S1), 5 mM MgCl_2_, 1 U μl^−1^ RNase inhibitor (APExBio), 50 mM Hepes pH 7.5, 200 mM NaCl, 25% glycerol, 0.5% Tween-20, and ∼2 ng RNA polymerase μl^−1^. The reaction mixtures were incubated for 1 h at 30°C, and stopped with formamide loading dye, denatured at 95 °C for 2 min and analyzed by 20% denaturing PAGE. The capped RNA cleavage products were visualized by phosphorimaging.

### Capped primer extension

To measure the extension and polyadenylation of mvRNA templates (Table S1) by the IAV RNA polymerase we followed the protocol described previously (33). Briefly, ∼2 ng RNA polymerase μl^−1^ was incubated for 10 min at 30°C in presence 1 mM dithiothreitol (DTT), 5 mM MgCl2, 1 U/μl RNase inhibitor (APExBio), 50 mM Hepes pH 7.5, 200 mM NaCl, 25% glycerol, 0.5% Tween-20 and 15 μM of balaxovir (Chemgenes). Next, the following components were added to the RNA polymerase mixture: 500 μM UTP, 500 μM ATP, 500 μM CTP, 500 μM GTP, ^32^P-labelled and capped 11 nucleotide-long RNA primer and 0.7 μM of mvRNA template (Table S1). The reaction mixtures were incubated for 1 h at 30°C and stopped with 4 μl formamide loading dye. Samples were subsequently denatured at 95 °C for 2 min and analyzed by 12% denaturing PAGE. Reaction products were visualized by phosphorimaging on a Tythoon scanner (GE Healthcare).

### T-loop and sequence analysis

The t-loop analysis was performed using a Python script as described previously. The distance between the entry and exit channels was set to 20 nt of the template sequence and the increment of the sliding window to 1 nt. The formation of secondary structures was measured by computing the ΔG between 6 nt upstream and 6 nt downstream of the location of the RNA polymerase. To analyze the length of U-tracts in mvRNA sequences, mvRNA sequences were retrieved from publicly available datasets described in (12).

### Statistical analysis

Experiments were repeated as indicated in each figure. Error bars in each figure indicate standard deviation. Data were plotted using Prism 10 (Graphpad) and analyzed using one-way ANOVA with Dunnet’s multiple comparisons test or two-way ANOVA where indicated in the figure legends.

## Acknowledgements

The authors would like to thank members of the te Velthuis lab for helpful discussions, in particular Michael Oade and Kimmie Sabsay. We thank Brandon Schweibenz and Joseph Macrotrigiano for sharing the purified RIG-I.

## Author contributions

EP, EE, and AJWtV conceived hypotheses. EP, KB, KR, and AJWtV performed experiments. AD and JY contributed resources. EP, KB, KR, and AJWtV analyzed biological data. EP and AJWtV performed structural and computational analyses. EP and AJWtV drafted the manuscript. All authors commented on and approved the manuscript.

## Data availability

All data are available from the authors upon request.

