## Supplementary material for "A two-step mechanism for RIG-I activation by influenza virus mvRNAs": Fig. S1

Promoter U-stretch

Segment 1 5'-AGUAGAAACAAGGUCGUUUUUA
Segment 2 5'-AGUAGAAACAAGGCAUUUUUUC
Segment 3 5'-AGUAGAAACAAGGUACUUUUUU
Segment 4 5'-AGUAGAAACAAGGGUGUUUUUC
Segment 5 5'-AGUAGAAACAAGGGUAUUUUUC
Segment 6 5'-AGUAGAAACAAGGAGUUUUUUG
Segment 7 5'-AGUAGAAACAAGGUAGUUUUUU
Segment 8 5'-AGUAGAAACAAGGGUGUUUUUU

**Figure S1. Sequence variation of the terminal vRNA sequence.** Sequence alignment based on A/WSN/1933 (H1N1) sequence.

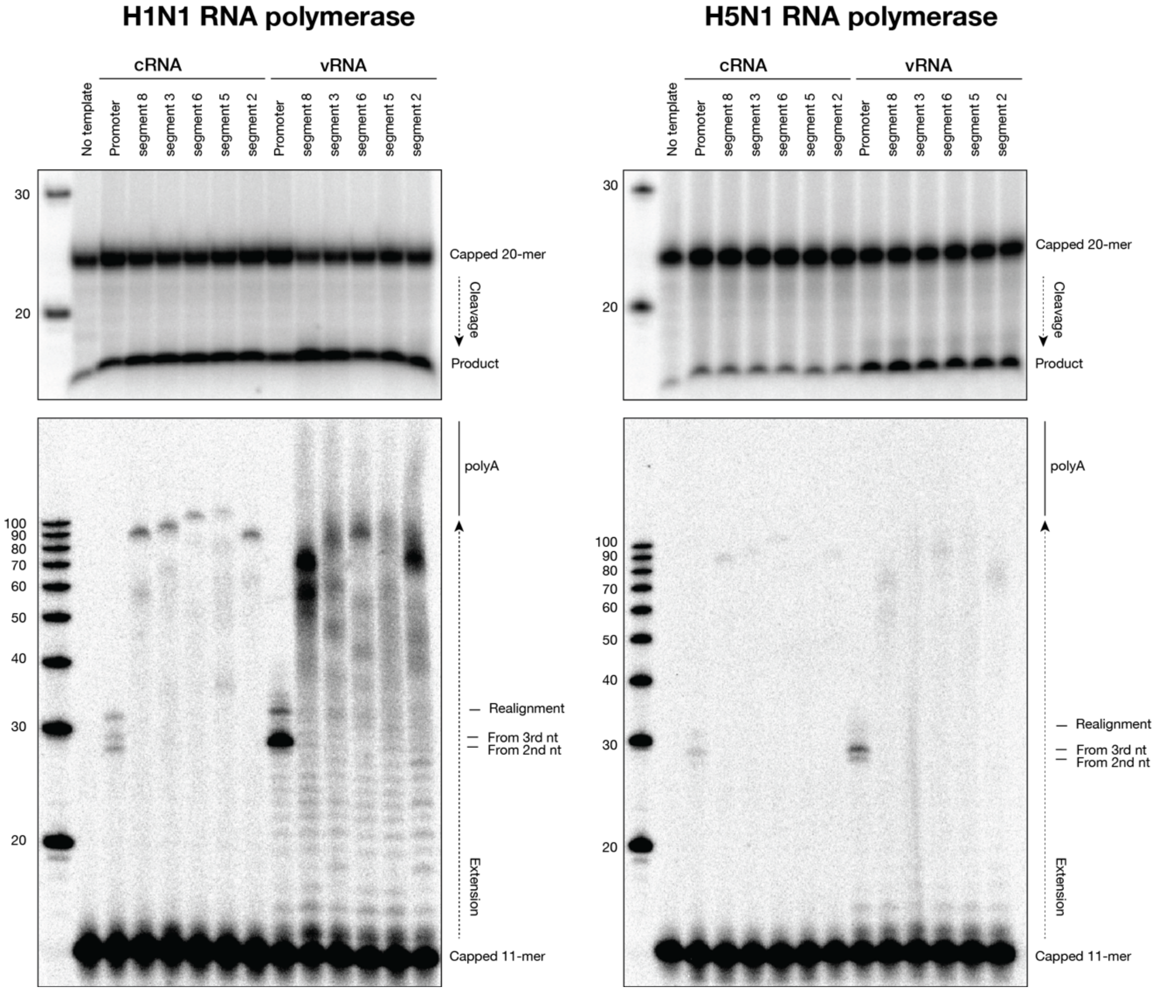


**Figure S2. Cap-snatching (top) and capped primer elongation activity (bottom) in vitro**. In the reactions shown, purified H1N1 or H5N1 IAV RNA polymerases were incubated with mvRNA templates derived from segments 8, 3, 6, 5 and 2. A representative image of two repeats is shown.

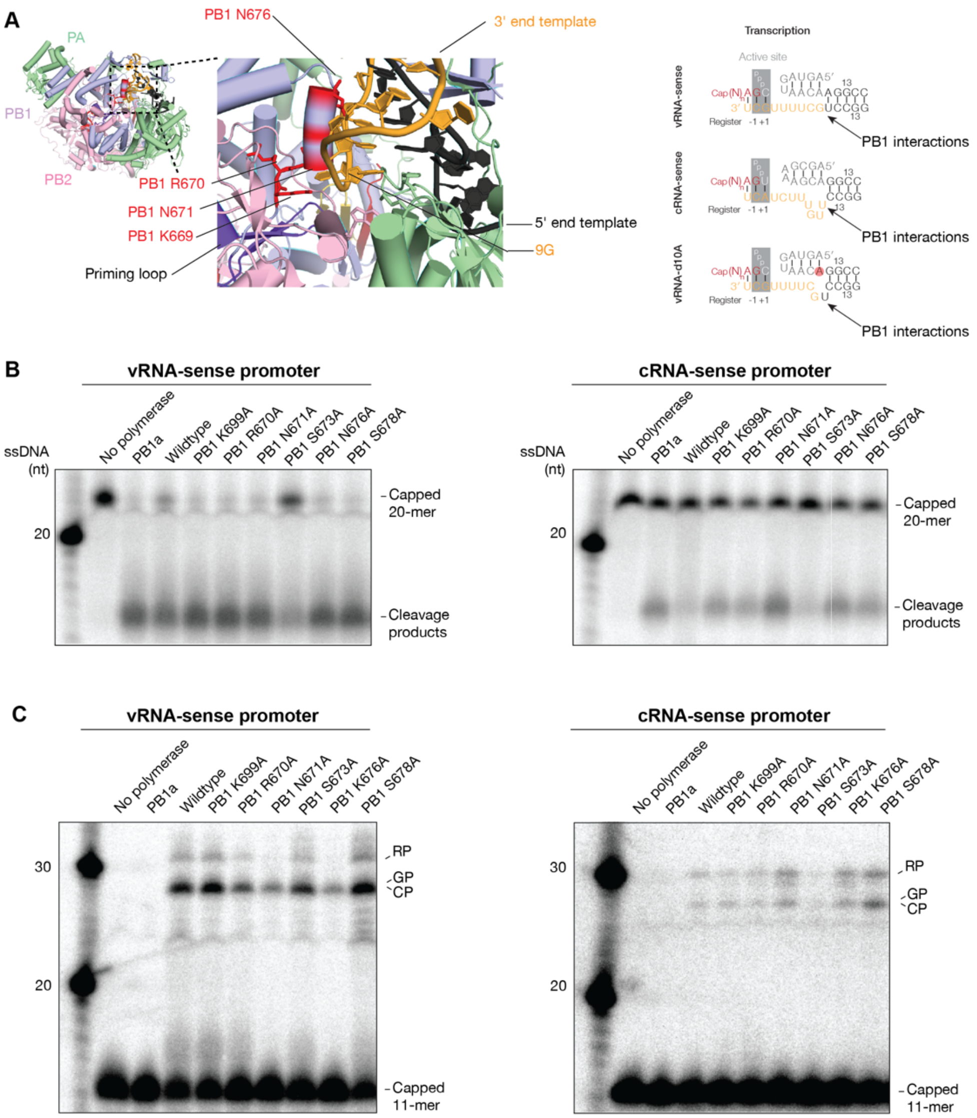


**Figure S3. Cap-snatching and capped primer elongation activity by PB1 mutants in vitro. A**) Structure of the IAV RNA polymerase (PDB 6qcs) with the positions of PB1 mutations indicated. The schematic on the right indicates the position where the PB1 helix interacts with the 3' end of the viral promoter sequence. **B and C**) Activity assay in vitro. In the reactions shown, purified WSN IAV RNA polymerase preparations were incubated with the IAV promoter 3' and 5' termini and either B) a radiolabeled capped 20-mer to measure endonuclease activity or C) a radiolabeled capped 11-mer and all four NTPs to measure capped primer extension. A representative image of two repeats is shown.

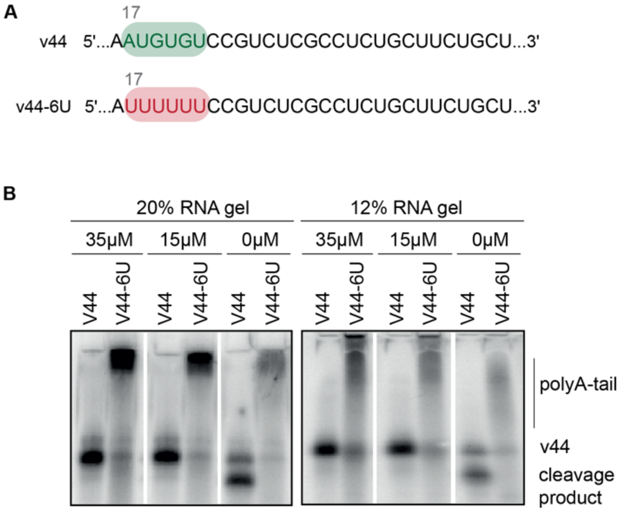

**Figure S4. Blocking cap-snatching using baloxavir in vitro**. **A**) RNA sequence of v44 and v44-6U based on Wandzik et al (reference 18) with a 6xU polyadenylation signal starting 17 nucleotides from the 5′ end. Position U17 is essential for polyadenylation and indicated in grey. **B**) In vitro transcription assay using the wildtype WNS RNA polymerase in presence of the RNA v44 or v44-6, with 0 µM, 15 µM or 35 µM of baloxavir (BAX). Products were separated by 20% or 12% denaturing PAGE in 7 M urea and detected by autoradiography. In the absence of BAX, extended products are cleaved by the IAV RNA polymerase, which confounds analysis of the data. A representative image is shown.

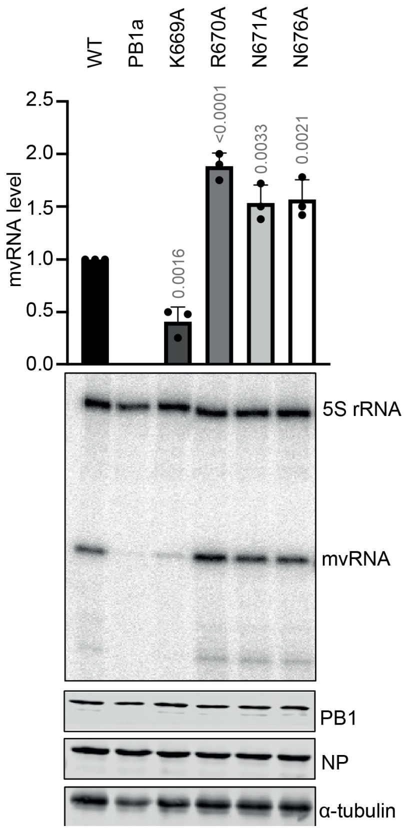


**Figure S5. Mutation of PB1 residues 669-676 does not lead to cvRNA synthesis in cells on an mvRNA template.** HEK293T cells were transfected with plasmids expressing the subunits of the IAV RNA polymerase and a plasmid expressing the NP71.2 mvRNA. Steady-state RNA levels were detected by primer extension. While shorter aberrant RNA products are visible below the mvRNA signal, no slower migrating products are observed for this mvRNA template. A representative image of three repeats is shown. Graph shows mean of three independent experiments. The p-values were calculated using one-way ANOVA and the error bars indicate standard deviation. Lower panels represent western blot protein expression controls.

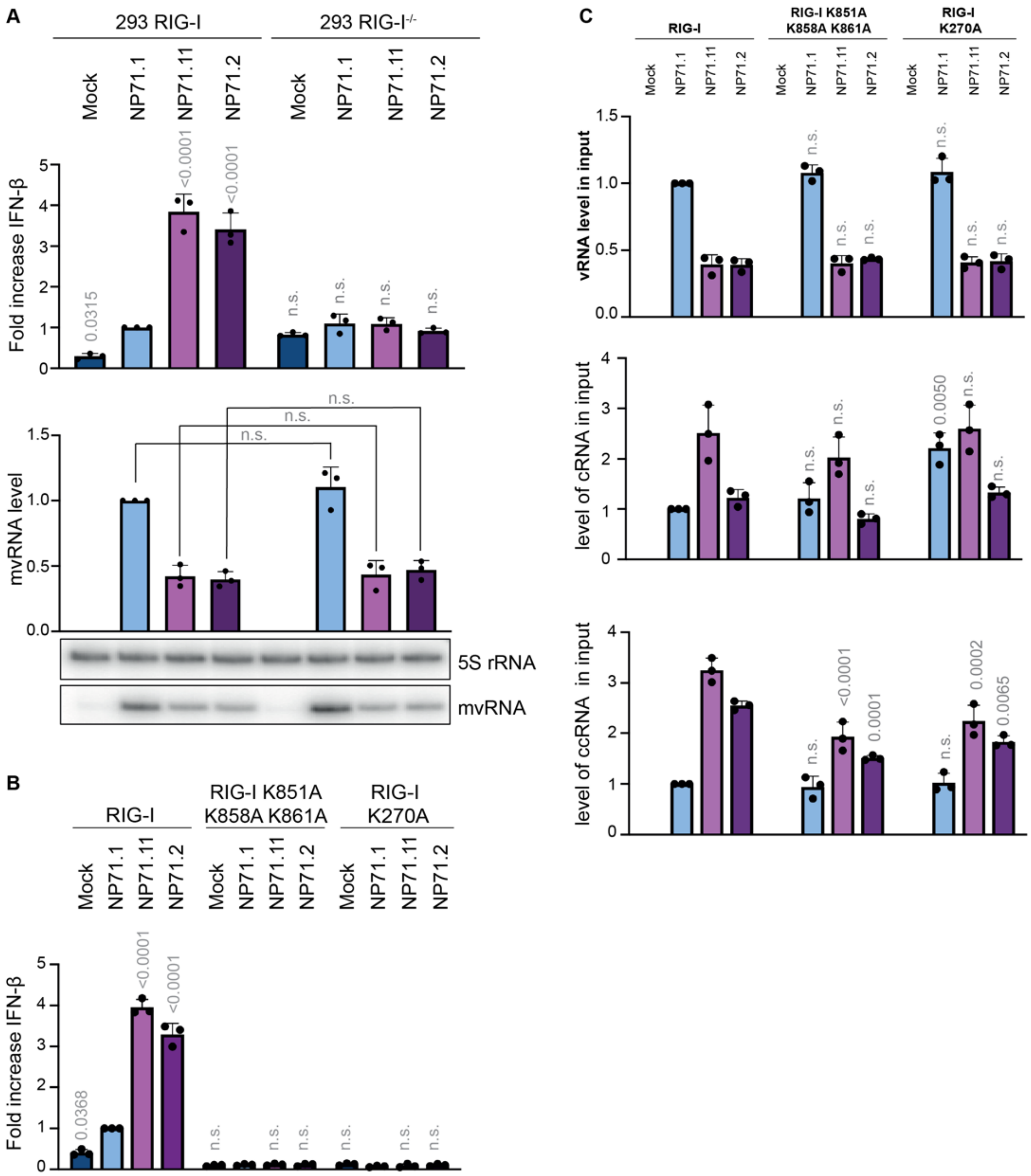


**Figure S6. Activation of innate immune signaling in RIG-I-dependent. A**) Analysis of IFN-β promoter activation in wildtype or RIG-I -/- HEK 293 cells using a luciferase reporter assay. Cells were transfected with segment 5 mvRNAs, the WSN IAV RNA polymerase, and NP. The luciferase signal was normalized to the *Renilla* signal and to the luciferase signal induced by the NP71.1 mvRNA in the wildtype cells. A representative image of three repeats is shown. **B**) IFN-β promoter activation in HEK 293T cells as analyzed using a luciferase reporter assay. Cells were transfected with the segment 5 mvRNAs, the WSN IAV RNA polymerase, NP, as well as c-myc RIG-I, RIG-I K851A-K858A-K861A, or RIG-I 270A. The luciferase signal was normalized to the *Renilla* signal and to the luciferase signal induced by the NP71.1 mvRNA. **C**) Input IAV RNA levels in HEK 293T cells transfected with segment 5 mvRNAs, the WSN IAV RNA polymerase, NP, and c-myc RIG-I, RIG-I K851A-K858A-K861A, or RIG-I 270A. In all graphs, the mean of three independent experiment is shown. The p-values were calculated using one-way ANOVA and the error bars indicate standard deviation.

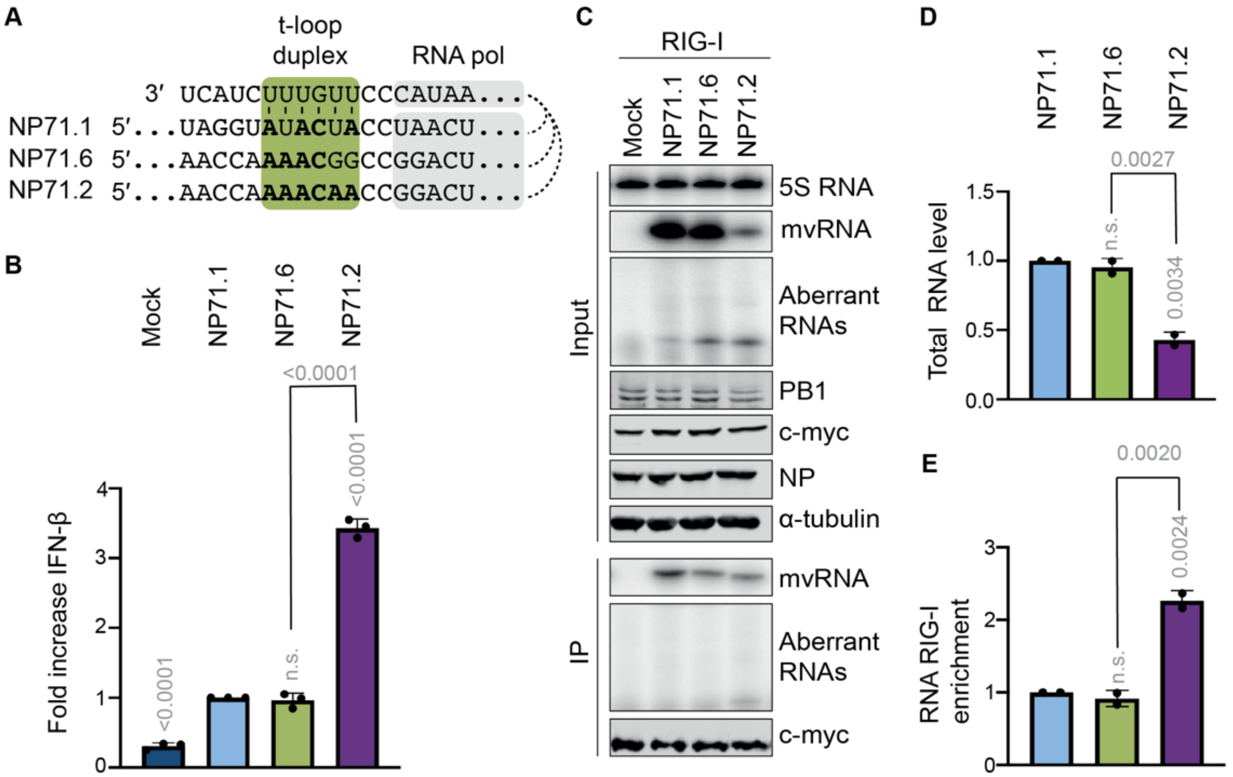


**Figure S7. Binding of mvRNA 71.6 to RIG-I. A**) Schematic showing point mutations introduced into the NP71.2 sequence to generate NP71.6. **B**) IFN-β promoter activation in HEK 293T cells transfected with the segment 5 mvRNAs, the WSN IAV RNA polymerase, and NP. Mean of three independent experiments is shown. The p-values were calculated using one-way ANOVA and the error bars indicate standard deviation. **C**) Primer extension analysis of RNA levels present in total RNA samples and RNA extracted after RIG-I immunoprecipitation. Representative images of two repeats are shown. **D**) Quantification of mvRNA level in total RNA relative to NP71.1. **E**) Quantification of mvRNA levels after immunoprecipitation relative to NP71.1. Mean of two independent experiments is shown. The p-values were calculated using one-way ANOVA and the error bars indicate standard deviation.

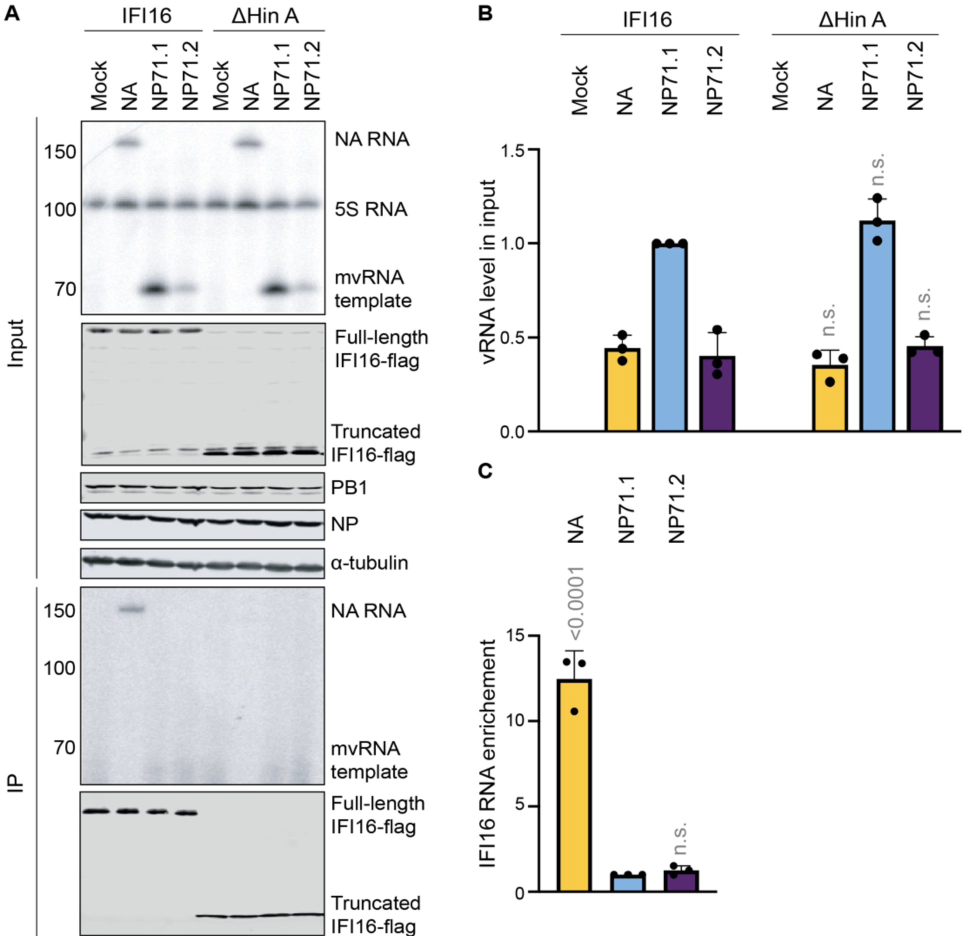


**Figure S8. IFI16 binding of IAV segment 6 and mvRNAs NP71.1 and NP71.2. A**) Immunoprecipitation of full-length IAV segment 6 or mvRNAs NP71.1 or NP71.2 by flag-tagged wildtype IFI16 or the IFI16 RNA binding mutant. A representative image of three repeats is shown. **B**) Input and **C**) immunoprecipitated RNA levels were measured using primer extension. In the graphs, the mean of three independent experiments is shown. The p-values were calculated using one-way ANOVA and the error bars indicate standard deviation.

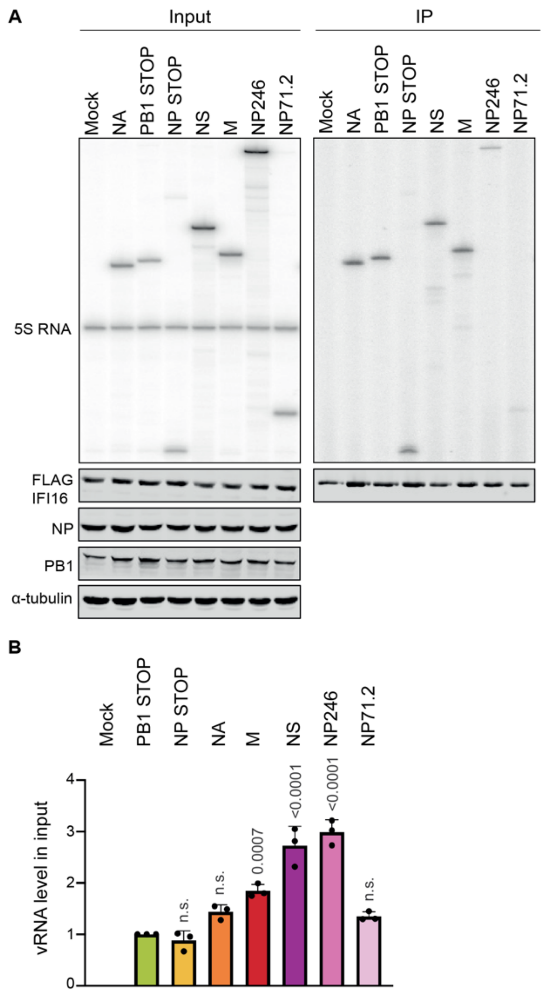


**Figure S9. IFI16 binding of IAV RNAs of different lengths. A**) Immunoprecipitation of full-length IAV segment 2, 5, 6, 7 or 8, a DelVG derived from segment 5, or mvRNA NP71.2 by flag-tagged wildtype IFI16. A representative image of three repeats is shown. **B**) Quantification of the IAV RNA levels in the input. The quantification of the immunoprecipitated RNAs is shown in main Fig 3.

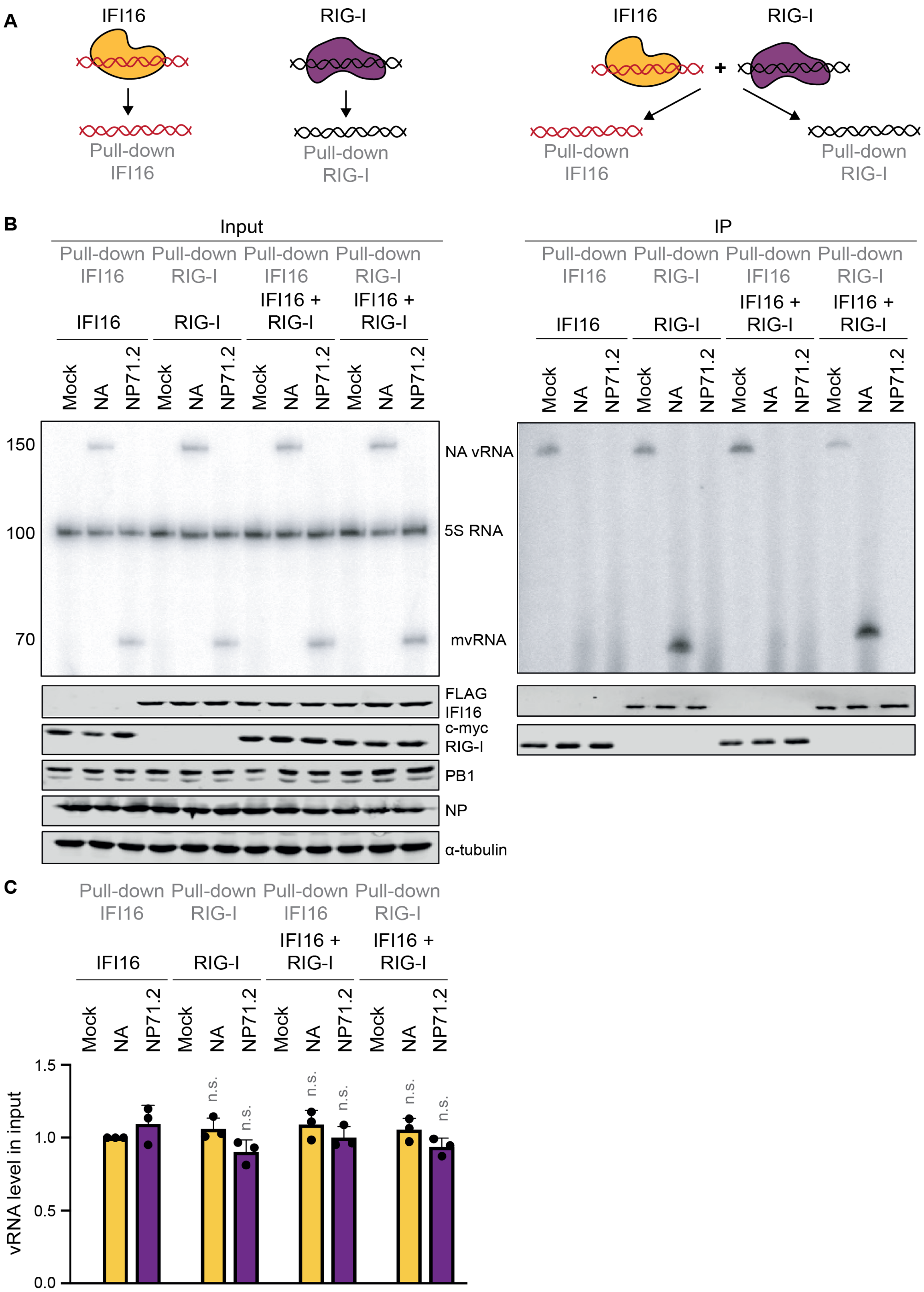


**Figure S10. IFI16 binding with or without RIG-I. A**) Schematic of four different immunoprecipitations to test the impact of RIG-I on IAV RNA binding. **B**) Immunoprecipitation of full-length IAV segment 6 or mvRNA NP71.2 by flag-tagged wildtype IFI16 or c-myc-tagged RIG-I either when the RNA binding proteins were expressed alone or when they were co-expressed. A representative image of three repeats is shown. **C**) Quantification of the input IAV RNA levels. The quantification of the immunoprecipitated RNAs is shown in main Fig 3.

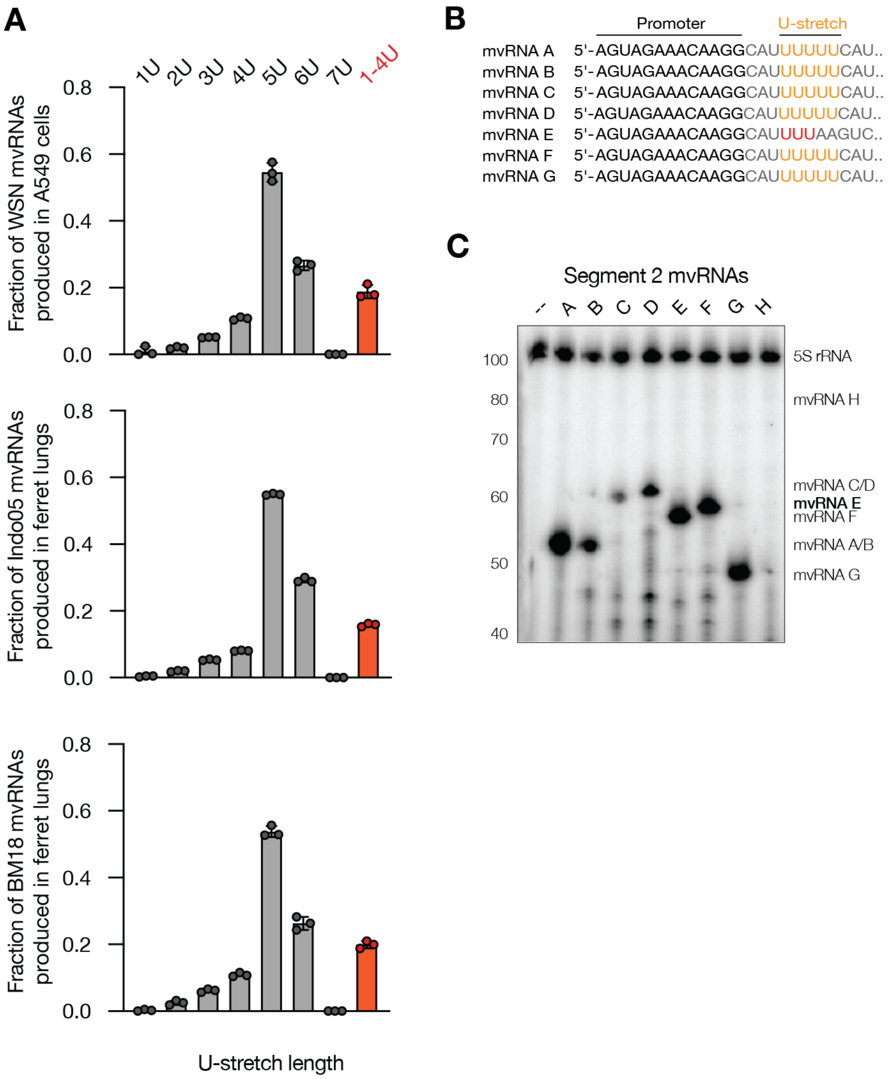


**Figure S11. ccRNAs production is upregulated on mvRNAs with truncated polyA signals. A**) Analysis of U-stretch length of mvRNAs detected in A549 cell infections with A/WSN/33 (H1N1) (WSN), in ferret lungs infected with A/Vietnam/1203/2004 (H5N1) (VN04), or in ferret lungs infected with A/Brevig Mission/1/1918 (H1N1) (BM18). **B**) Alignment of the U-tracts of cloned segment 2 mvRNAs. **C**) Primer extension analysis of the negative sense RNA molecules produced during mvRNA replication.

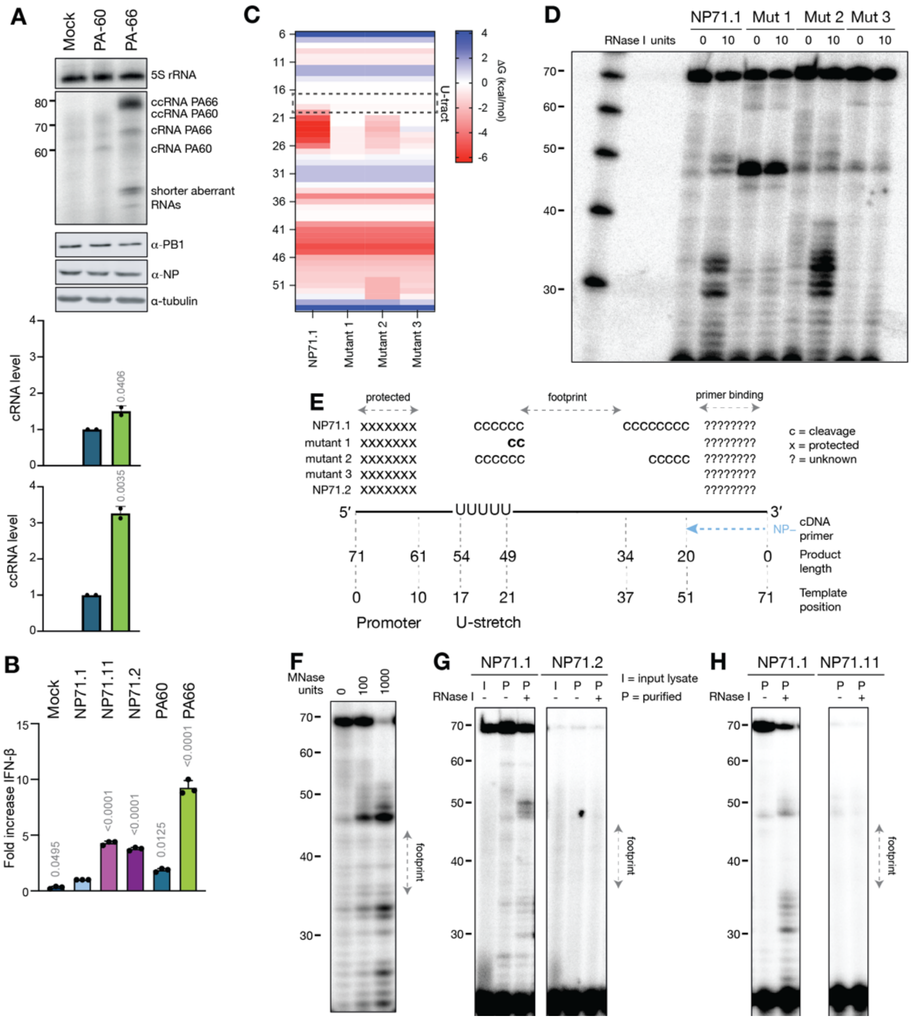


**Figure S12. Foot printing analysis of various mvRNAs.** **A**) Steady state ccRNA, cRNA and 5S rRNA levels measured 24 hours post-transfection by primer extension. PB1, NP, and tubulin expression were analyzed by western blot. Quantified steady state RNA levels following are shown in the bottom graphs. **B**) Innate immune activation by IAV mvRNAs in HEK 293T cells. Cells were transfected with plasmids encoding the RNA polymerase subunits, NP, a *Renilla* luciferase transfection control, a firefly-based IFN-$\beta$ reporter, and the mvRNAs indicated. Twenty-four hours post-transfection, the luciferase signal was measured. **C**) Heatmap showing the stability of t-loops in the NP71.1 mvRNA and mutants 1, 2 and 3. **D**) Analysis of NP71.1 footprint or NP71.1 containing single (Mut 1 or 2) or double (Mut 3) mutations to destabilize the t-loop in NP71.1. **C**) Summary of segment 5 mvRNA foot printing data shown in Fig. S12. **F**) Analysis of NP71.1 footprint using different amounts of MNase. **G**) Analysis of NP71.1, NP71.2 and footprints. **H**) Analysis of NP71.1, NP71.11 and footprints. For all experimental panels, a representative image is shown. In panels A and B, graphs show mean of 2 or 3 independent experiments. Error bars indicate standard deviation and p-values were determined using one-way ANOVA.

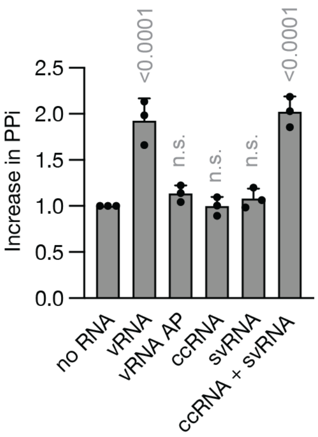


**Figure S13. RIG-I ATPase activity as measured using thin layer chromatography.** As negative control, mvRNA in vitro transcripts were treated with alkaline phosphatase (AP). Error bars represent standard deviation. P-values were calculated using one-way ANOVA relative to the no RNA control.

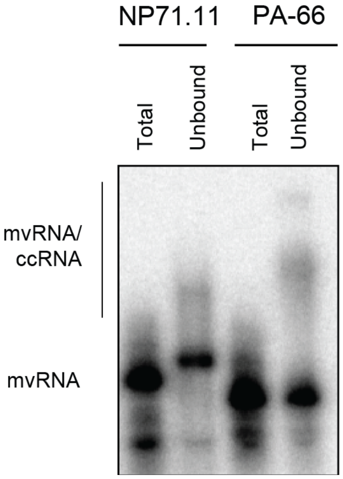


**Figure S14. Non-denaturing PAGE analysis of ccRNA-released fraction.** The unbound fractions of NP71.11 and PA-66 ccRNA-treated reactions were analyzed by 6% non-denaturing PAGE. For comparison, RNA-polymerase-bound mvRNAs samples prior to treatment with ccRNA were also analyzed. The gel-shifts relative to the mvRNA control lanes, which likely represent mvRNA/ccRNA duplexes, is indicated.

**Table S1**

| Template Name | Sequence of the RNA templates (5′ to 3′) |
|---|---|
| NP STOP | WSN NP full-length STOP (Start codon ATG mutated in stop codon TCG) |
| NA | WSN NA full length |
| PB1 STOP | WSN PB1 full length STOP (Start codon ATG mutated in stop codon TCG) |
| M | WSN M Full length |
| NP246 | AGUAGAAACAAGGGUAUUUUUCUUUAAUUGUCGUACUCCUCUGCAUUGUCUCCGAAG AAAUAAGAUCCUUCAUUACUCAUGUCAAAGGAGGGCACGAUCGGGCUCGUUGCCUUU UCGUCCGAGAGCUCGAAGACUCCCCGCCCCUGGAAAGACACUAGUCUCCAUCUGUUC GUAAGAUCGUUUGGUGCCUUUGGUCGCCAUGAUUUCGAUGUCACUCUGUACUAGUC UACCCUGCUUUUGCU |
| NP71.1 | AGUAGAAACAAGGGUAUUUUUCUUUACUAGUUAGGUAGUAUACCUAGUAACUAGUCU ACCCUGCUUUUGCU |
| NP71.11 | AGUAGAAACAAGGGUAUUUUUCUUUACUAGUGGCAGCAAAAGCACCCAUACUAGUCU ACCCUGCUUUUGCU |
| NP71.2 | AGCAAAAGCAGGGUAGACUAGUGGCAACCAAAACAACCGGACUAGUAAAGAAAAAUAC CCUUGUUUCUACU |
| NP71.6 | AGCAAAAGCAGGGUAGACUAGUGGCAACCAAAACGGCCGGACUAGUAAAGAAAAAUA CCCUUGUUUCUACU |
| PA-60 | AGUAGAAACAAGGUACUUUUUUGGACAGUAUGCCAUUUUGAAUCAGUACCUGCUUUC GCU |
| PA-66 | AGUAGAAACAAGGUACUUUUUUGGACAGUAUGGAUAGCACAUUUUGAAUCAGUACCU GCUUUCGCU |
| PB1-A | AGUAGAAACAAGGCAUUUUUUCAUGAAAUCCAUUCAAAUGGUUUGCCUGCUUUCGCU |
| PB1-B | AGUAGAAACAAGGCAUUUUUUCAUGAAGGACAUUCAAAUGGUUUGCCUGCUUUCGCU |
| PB1-C | AGUAGAAACAAGGCAUUUUUUCAUGAAGGACAAGCUAAACAUUCAAAUGGUUUGCCU GCUUUCGCU |
| PB1-D | AGUAGAAACAAGGCAUUUUUUCAUGAAGGACAAGCUAAAUCAUUCAAAUGGUUUGCC UGCUUUCGCU |
| PB1-E | AGUAGAAACAAGGCAUUUUAAGUCGGAUUGACAUCCAUUCAAAUGGUUUGCCUGCUU UCGCU |
| PB1-F | AGUAGAAACAAGGCAUUUUUUCAGUCGGAUUGACAUCCAUUCAAAUGGUUUGCCUGC UUUCGCU |
| PB1-G | AGUAGAAACAAGGCAUUUUUUCAUGCAUUCAAAUGGUUUGCCUGCUUUCGCU |
| PB1-H | AGUAGAAACAAGGCAUUUUUUCAUGAAGGACAAGCUAAAUUCAGUUUGCCUGCUUUC GCU |
| NP71.1 mut1 | AGUAGAAACAAGCCUAUUUUUCUUUACUAGUUAGGUAGUAUACCUAGUAACUAGUCU ACCCUGCUUUUGCU |
| NP71.1 mut2 | AGUAGAAACAAGGGUAUUUUUCUUUACUAGUUAGGUAGUAUAGCUAGUAACUAGUCU ACCCUGCUUUUGCU |
| NP71.1 mut3 | AGUAGAAACAAGGGUAUUUUUCUUUACUAGUUAGGUAGUAUAGGUAGUAACUAGUCU ACCCUGCUUUUGCU |
| cRNA 3’ | GGCCUUGUUUCUACU |
| cRNA 5’ | AGCGAAAGCAGGCC |
| vRNA 3’ | GGCCUGCUUUUGCU |
| vRNA 5’ | AGUAGUAACAAGGCC |

**Table S2**: Sequences of DNA oligonucleotides used for mutagenesis of the pcDNA3-PB1 expression plasmid and the NP71.1 t-loop located upstream of the U-tract.

| Primer | Sequence of the DNA oligonucleotide (5′ to 3′) |
|---|---|
| pcDNA3-PB1 K669A Fwd | CACTCCTGGATCCCCGCAAGAAATCGATCCATC |
| pcDNA3-PB1 K669A Rev | GATGGATCGATTTCTTGCGGGGATCCAGGAGTG |
| pcDNA3-PB1 R670A Fwd | CTCCTGGATCCCCAAAGCAAATCGATCCATCTTG |
| pcDNA3-PB1 R670A Rev | CAAGATGGATCGATTTGCTTTGGGGATCCAGGAG |
| pcDNA3-PB1 N671A Fwd | GGATCCCCAAAAGAGCTCGATCCATCTTGAATACAAGC |
| pcDNA3-PB1 N671A Rev | GCTTGTATTCAAGATGGATCGAGCTCTTTTGGGGATCC |
| pcDNA3-PB1 S673A Fwd | GGATCCCCAAAAGAAATCGAGCTATCTTGAATACAAGCC |
| pcDNA3-PB1 S673A Rev | GGCTTGTATTCAAGATAGCTCGATTTCTTTTGGGGATCC |
| pcDNA3-PB1 N676A Fwd | GGATCCCCAAAAGAGCTCGATCCATCTTGAATAC |
| pcDNA3-PB1 N676A Rev | GTATTCAAGATGGATCGAGCTCTTTTGGGGATCC |
| pcDNA3-PB1 S678A Fwd | CGATCCATCTTGAATACAGCACAAAGAGGAATACTTG |
| pcDNA3-PB1 S678A Rev | CAAGTATTCCTCTTTGGCTTGTATTCAAGATGGATCG |
| Mut 1 G13C G14C Fwd | TAGAAACAAGccTATTTTTCTTTACTAGTTAGGTAG |
| Mut 1G13C G14C Rev | CTAATAACCCGGCGG |
| Mut 2 G43C Fwd | AGGTAGTATAgCTAGTAACTAG |
| Mut 2 G43C Rev | AACTAGTAAAGAAAAATACCC |
| Mut 3 G43C G44C Fwd | AGGTAGTATAggTAGTAACTAGTCTACCCTG |
| Mut 3 G43C G44C Rev | AACTAGTAAAGAAAAATACCC |

**Table S3: DNA oligonucleotides used for primer extension.**

| Primer name | Target RNA | DNA oligonucleotide (5' to 3') |
|---|---|---|
| NP- | Non-full-length NP vRNA and aberrant products | AGCAAAAGCAGGGTAGACTAGT |
| NP+ | Non-full-length NP mRNA | ACTAGTCTACCCTGCTTTTGC |
| NP5' | Non-full-length NP cRNA | AGTAGAAAACAAGGGTATTTTTC |
| PA_PEplus2 | Non-full-length PA cRNA | AGTAGAAACAAGGTACTTTTTTGGACAGTATGG |
| NA1280 | Full length NA vRNA | TGGACTAGTGGGAGCATCAT |
| NA160 | Full length NA cRNA and mRNA | TCCAGTATGGTTTTGATTTCC |
| PB1vRNA | Full length PB1 vRNA | TGATTTCGAATCTGGAAGGA |
| PB1c/mRNA | Full length PB1 cRNA and mRNA | TCCATGGTGTATCCTGTTCC |
| MvRNA | Full length M vRNA | GAAAAGAGGGCCTTCTACGG |
| Mc/mRNA | Full length M cRNA and mRNA | AGCCATTCCATGAGAACCTC |
| NP149- | Full length NP vRNA | ATTTCTTCGGAGACAATGCAG |
| NP149+ | Full length NP cRNA and mRNA | TAAGATCGTTTGGTGCCTTTG |
| NSvRNA | Full length NS vRNA | TGATTGAAGAAGTGAGACACAG |
| NSc/mRNA | Full length NS cRNA and mRNA | CGCTCCACTATTTGCTTTCC |
| 5S_100 | 5S sRNA; loading control | TCCCAGGCGGTCTCCCATCC |
| 5S_62 | 5S sRNA; loading control | ACCCTGCTTAGCTTCCGAGA |

**Table S4: Antibody used for western blot.**

| Primary antibody | | |
|---|---|---|
| NP | Rabbit, GTX125989, GeneTex | 1:4000 |
| PB1 | Rabbit, GTX125923, GeneTex | 1:1000 |
| PA | Rabbit, GTX125932, GeneTex | 1:1000 |
| $\gamma$-tubulin | Rat, MCA77G, Bio-Rad | 1:5000 |
| MitoTracker | Mouse, AB92824, Abcam | 1:1000 |
| Histone H3 | Rabbit, AB1791, Abcam | 1:1000 |
| ANP32A | Rabbit, 15491S, Cell signaling | 1:1000 |
| NA H1N1 | Rabbit, GTX125974, GeneTex | 1:1000 |
| HA H1N1 | Rabbit, GTX127357, GeneTex | 1:500 |

| | | |
|---|---|---|
| RIG-I | Mouse, SC-376845, Santacruz | 1:1000 |
| EQKLISEEDL c-myc tag | Mouse, M4439, Sigma | 1:1000 |
| Phospho-IRF3 | Rabbit, 29047S, Cell signaling | 1:1000 |
| DYKDDDDK Flag tag | Rabbit, PA1-984B, Invitrogen | 1:1000 |
| Phospho-STAT1 (Tyr701) | Rabbit, MA5-15071, Thermofischer | 1:500 |
| STAT1 | Mouse, MA1-037, Thermofischer | 1:300 |
| Secondary antibody | | |
| IRDye 680 goat anti-mouse | 926-68020, LI-COR | 1:10000 |
| IRDye 800 goat anti-mouse | 926-32210, LI-COR) | 1:10000 |
| IRDye 680 goat anti-rat | 926-68076, LI-COR | 1:10000 |
| IRDye 680 goat anti-rabbit | 926-68071, LI-COR | 1:10000 |
| IRDye 800 donkey anti-rabbit | 926-32213, LI-COR | 1:10000 |
